# Median raphe input to dorsal CA1 shapes VIP interneuron recruitment and novelty-guided spatial memory

**DOI:** 10.64898/2026.09.17.752379

**Authors:** Xiao Luo, Risna Radhakrishnan, Alexandre Guet-McCreight, Dimitry Topolnik, Aurélie Bergeron, Vandana Dulam, Louis-David Coulombe, Megan Besner, Justine Fortin-Houde, Judy Davies, Félix Michaud, Bénédicte Amilhon, Lisa Topolnik

## Abstract

Vasoactive intestinal peptide–expressing interneurons (VIP-INs) gate hippocampal inhibition during novel experience, but the long-range signals that engage these cells remain poorly understood. Here we identify median raphe (MnR) projections as a brainstem pathway that tunes dorsal CA1 VIP-IN recruitment through coordinated glutamatergic and serotonergic mechanisms. Anatomical mapping and optogenetic recordings showed that MnR axons innervate multiple VIP-IN subtypes, while transcriptomic and pharmacological analyses revealed fast glutamatergic excitation together with serotonin receptor–dependent modulation of synaptic and intrinsic responsiveness. In vivo calcium imaging showed that novelty preferentially recruited a speed-coupled VIP-IN ensemble, and inhibition of MnR input selectively reduced the magnitude of this response. A hippocampal circuit model linked this pathway to dendritic disinhibition and place-cell recruitment. Behaviorally, inhibition of MnR input preserved exploratory engagement but disrupted the organization of spatial sampling and impaired object-location memory. Thus, MnR input organizes hippocampal disinhibition to support novelty-guided exploration and memory encoding.

## Introduction

Novel experience requires hippocampal circuits to maintain representational stability while transiently opening a window for new encoding. This balance between stability and plasticity is unlikely to arise from a global increase in excitation. Instead, accumulating evidence indicates that novelty rapidly reorganizes local inhibition, creating a permissive circuit state in which selected inputs can reshape hippocampal activity without destabilizing the network as a whole. Early intracellular recordings in freely moving rats provided some of the first evidence that exposure to a novel environment is associated with a transient reduction in hippocampal inhibition (Wilson and McNaughton, 1993; Nitz and McNaughton, 2004). More recent two-photon calcium imaging from genetically targeted interneuron populations further showed that distinct inhibitory classes, including parvalbumin (PV) and somatostatin (SST) -expressing interneurons, decrease their activity during novel-environment exploration (Arriaga and Han, 2019; Geiller et al., 2020; Hainmueller et al., 2024). Thus, novelty appears to engage a structured reconfiguration of inhibition, but the interneuron motifs and long-range pathways that initiate this state remain incompletely understood.

Vasoactive intestinal peptide–expressing interneurons (VIP-INs) are well positioned to coordinate such inhibitory reconfiguration (Kullander and Topolnik, 2022). Across cortical circuits, VIP-INs preferentially innervate other interneurons, including calbindin-, SST-, and PV-expressing cells, and are recognized as key regulators of inhibition (Acsady et al., 1996; Dalezios et al., 2002; Staiger et al., 2004; David et al., 2007; Guet-McCreight et al., 2020). Consistent with this organization, hippocampal CA1 VIP-INs are largely embedded in disinhibitory motifs (Francavilla et al., 2015; Kullander and Topolnik, 2022). With the exception of VIP basket cells, they predominantly contact other interneurons, including oriens-lacunosum moleculare (OLM) cells, bistratified cells, and basket cells (Tyan et al., 2014; Francavilla et al., 2018; Topolnik and Tamboli, 2022).

Hippocampal VIP-INs, however, are not a uniform population. They comprise multiple subtypes with distinct molecular, morphological, physiological, and connectivity profiles, including cholecystokinin-coexpressing VIP basket cells (VIP/CCK-BCs), calretinin-negative interneuron-specific type 2 cells, calretinin-positive interneuron-specific type 3 cells (VIP/CR-IS-3), muscarinic receptor 2-coexpressing long-range projecting VIP cells (VIP/M2R-LRPs), and additional VIP-IN populations distributed across CA1 layers (Acsady et al., 1996; Tyan et al., 2014; Francavilla et al., 2018; Harris et al., 2018; Geiller et al., 2020; Luo et al., 2019; 2020; Turi et al., 2019). This diversity suggests that VIP-INs do not provide a single disinhibitory signal. Rather, distinct VIP-IN subtypes may be recruited by different inputs and behavioral states to regulate specific inhibitory motifs.

Consistent with this view, hippocampal VIP-INs have been implicated in several behaviorally relevant computations, including goal-directed spatial learning (Magnin et al., 2019; Turi et al., 2019), lateral entorhinal cortex–driven dendritic gating (Bilash et al., 2023), CA2-dependent social memory (Leroy et al., 2022), and CA1 novelty detection (Tamboli et al., 2024; Neubrandt et al., 2025). In freely moving mice, entry into a novel context evokes a rapid and transient increase in dorsal CA1 (dCA1) VIP-IN activity, followed by pyramidal cell activation (Tamboli et al., 2024). dCA1 VIP-INs also respond to object novelty, including the introduction of new objects and the displacement of familiar objects to new locations. Moreover, optogenetic inhibition of VIP-INs during object sampling impairs subsequent object recognition memory, supporting a role for VIP-INs in enabling hippocampal circuits to encode novel information (Tamboli et al., 2024). These findings raise a central question: which inputs recruit dCA1 VIP-INs during novel experience, and do they shape the way novelty is explored and encoded?

Monosynaptic rabies tracing has shown that dCA1 VIP-INs receive input from multiple cortical and subcortical regions implicated in hippocampal state regulation, novelty processing, and learning, including the medial septum, ventral tegmental area, and median raphe (MnR) nucleus (Turi et al., 2019). These afferent pathways are likely to engage VIP-INs in different behavioral contexts and through distinct signaling mechanisms. The MnR provides a particularly compelling candidate input because it combines fast glutamatergic excitatory transmission with serotonergic neuromodulation, two signaling modes well suited to recruit and tune interneuron activity during rapid state transitions (Kocsis et al., 2006; Jackson et al., 2008; Wang et al., 2015; Domonkos et al., 2016). Although MnR circuits have been prominently linked to anxiety- and stress-related behaviors, some MnR neurons also fire during sensory stimulation and are coupled to hippocampal theta oscillations (Viana Di Prisco et al., 2002; Domonkos et al., 2016; Huang et al., 2022), placing MnR input in a position to coordinate exploratory state with hippocampal spatial processing.

The MnR contains intermingled serotonergic (5-hydroxytryptamine; 5-HT), vesicular glutamate transporter 3-expressing (VGLUT3), dual 5-HT/VGLUT3, GABAergic, and additional VGLUT2-expressing neurons (Sos et al., 2017; Ren et al., 2019; Szonyi et al., 2019). Ascending MnR projections to hippocampal CA1 are prominently glutamatergic and VGLUT3-positive, and can exert fast excitatory control over local circuits involved in hippocampal learning (Jackson et al., 2009; Varga et al., 2009; Szonyi et al., 2016; Xu et al., 2021; Fortin-Houde et al., 2023). In parallel, MnR 5-HT projections innervate diverse inhibitory interneuron classes and can contribute to network synchronization, excitation/inhibition balance, and memory-related hippocampal dynamics (Vertes, 1981, 1999; Kohler and Steinbusch, 1982; McMahon and Kauer, 1997; Gulyas et al., 1999; Nitz and McNaughton, 1999; Varga et al., 2009; Wang et al., 2015; Fernandez et al., 2017; Teixeira et al., 2018). In vivo two-photon imaging has further revealed functionally distinct streams of MnR 5-HT input associated with reward and locomotion (Luchetti et al., 2020). Together, these observations position the MnR as a versatile brainstem node capable of shaping hippocampal circuits through multiple transmitter systems.

Several features make MnR projections especially relevant for CA1 VIP-INs. Prior anatomical tracing in rat hippocampus indicated that MnR 5-HT fibers can contact VIP-INs (Papp et al., 1999). In addition, VIP-INs belong to the broader class of 5-HT3a receptor–expressing interneurons, characterized by strong expression of this fast ionotropic receptor in both neocortex and hippocampus (Lee et al., 2010; Rudy et al., 2011). However, whether MnR inputs target specific CA1 VIP-IN subtypes, how these transmitter systems regulate VIP-IN responsiveness, and whether this pathway shapes VIP-IN activity and novelty-guided exploration remain unresolved.

Here, we addressed these questions in dCA1 using anatomical tracing, optogenetic circuit mapping, patch-clamp recordings, patch-seq, simultaneous in vivo calcium imaging and optogenetic intervention, computational modeling and behavioral analysis. We show that MnR inputs modulate CA1 VIP-INs through subtype-specific connectivity motifs and complementary glutamatergic and serotonergic mechanisms. During novel-context exploration, VIP-INs form functionally distinct ensembles that are differentially related to novelty and locomotion, with MnR input selectively amplifying a VIP-IN population enriched for positively velocity-modulated VIP/CR-IS-3 cells. Inhibiting MnR–dCA1 projections leaves baseline exploration largely intact but reduces VIP-IN recruitment, alters novelty-guided spatial sampling, and impairs object-location memory. These findings identify a brainstem–hippocampal pathway that organizes VIP-IN ensemble activity and supports exploratory structure and spatial encoding during novelty-guided exploration.

## Results

### Median raphe projections excite distinct VIP interneuron subtypes

We first asked how MnR projections are organized with respect to the diverse VIP-IN population in dCA1. To visualize MnR axons and enable circuit mapping, we expressed channelrhodopsin-2 (ChR2) in the MnR of Vip-IRES-Cre-Ai9 mice using AAV5-hSyn-ChR2(H134R)-EYFP (Fig. 1a). Immunohistochemical analysis confirmed viral expression in both serotonergic and VGLUT3-expressing MnR neurons (Supplementary Fig. 1a, b), whereas tdTomato expression in dCA1 showed high specificity and sensitivity for VIP-INs (Supplementary Fig. 1c, d).

**Figure 1.**
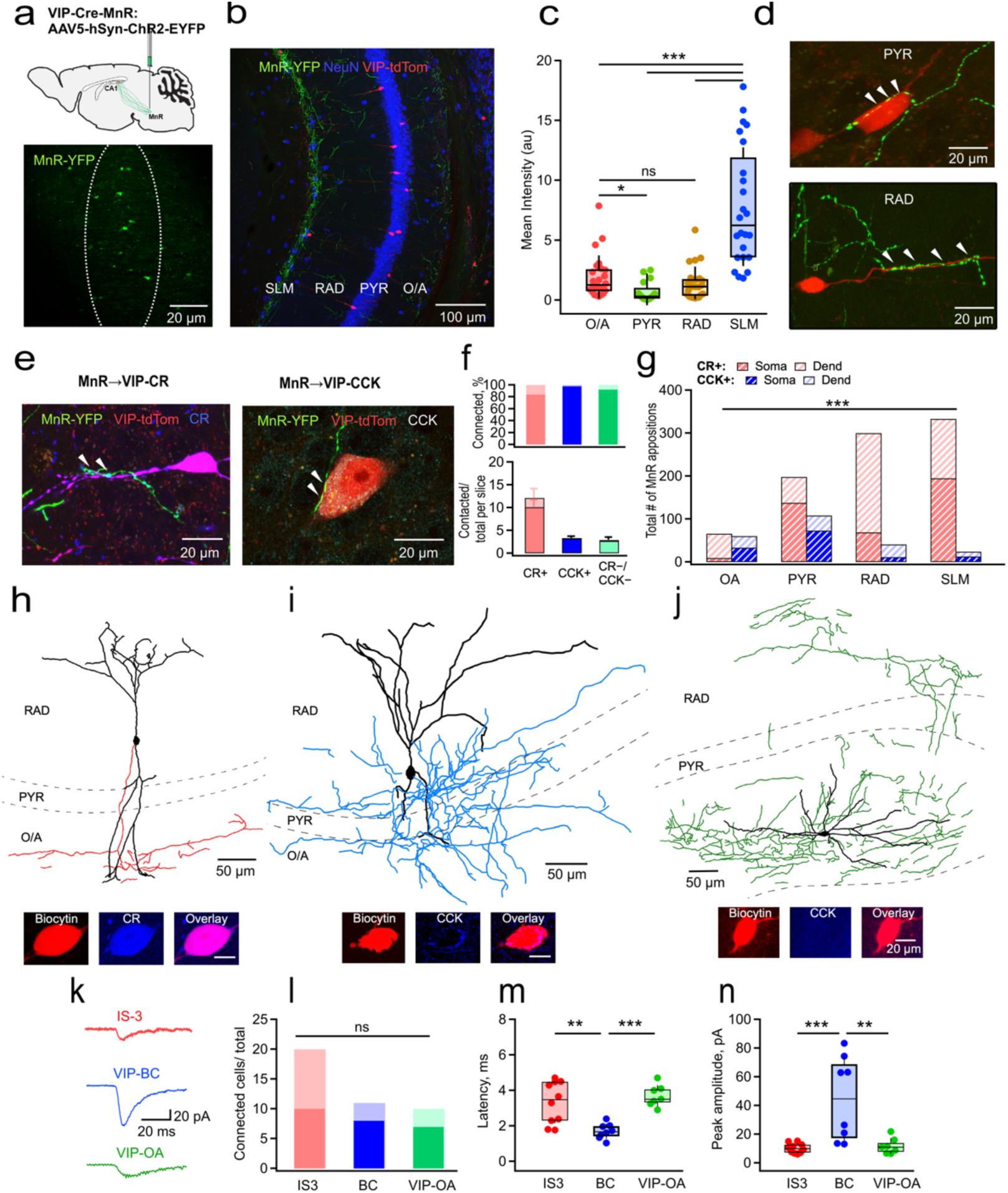
Anatomical and functional mapping of MnR inputs to CA1 VIP interneurons. **a,** Viral strategy for expression of ChR2–eYFP in the MnR of Vip-IRES-Cre-Ai9 mice and representative MnR expression. **b,** Representative dCA1 image showing MnR axons, NeuN-defined CA1 layers, and tdTomato-expressing VIP-INs. **c,** MnR-eYFP signal across CA1 layers (H(3) = 49.24, p = 1.16 x 10^-10^; Kruskal-Wallis test; n = 24 sections from 4 mice, 2 male/2 female; see Supplementary Table 1 for full statistical details). **d,e,** Representative putative MnR axonal appositions onto VIP-IN somata and dendrites (**d**) and onto CR+ or CCK+ VIP-INs (**e**). Arrows indicate putative appositions. **f,** Number and fraction of CR+, CCK+, and CR−/CCK− VIP-INs with putative MnR appositions (n = 172/205 CR+, 45/46 CCK+, and 35/38 CR−/CCK− cells from 17 sections and 3 mice). **g,** Aggregate distribution of putative somatic and dendritic appositions by VIP-IN subtype and CA1 layer (H(7) = 27.29, pv= 0.0003; Kruskal-Wallis test; n = 172 CR+ cells, 45 CCK+ cells; 17 sections from 3 mice, 2 male/1 female; see Supplementary Table 1 for full statistical details). **h–j,** Morphological reconstructions and post hoc immunolabeling of a VIP/CR-IS-3 cell (**h**), VIP/CCK basket cell (**i**), and VIP-OA cell (**j**). Soma and dendrites are shown in black and axons are shown in red (IS-3), blue (BC) or green (VIP-OA). **k,** Representative light-evoked MnR EPSCs. **l,** Functional connection probability across VIP-IN subtypes (10/20 IS-3 cells from 9 mice, 8/11 VIP/CCK-BCs from 6 mice, and 7/10 VIP-OA cells from 5 mice; p = 0.418; two-sided Fisher-Freeman-Halton test; see Supplementary Table 1 for full statistical details). **m,n,** Latency (**m,** F(2,22) = 12.33, p = 0.0002, one-way Welch ANOVA) and peak amplitude (**n,** F(2,22) = 11.09, p = 0.00026, one-way Welch ANOVA) of connected responses (n = 10, 8, and 7 cells, respectively, from 20 mice). Box plots show median and interquartile range; points represent cells. Full statistical details are provided in Supplementary Table 1. *\*\*p < 0.01, ***p < 0.001;* ns, not significant.

Consistent with prior anatomical studies (Freund et al., 1990; Papp et al., 1999; Szonyi et al., 2016; Fortin-Houde et al., 2023), MnR axons were distributed across all CA1 layers, with the highest density in stratum lacunosum-moleculare (SLM), followed by stratum radiatum (RAD), oriens/alveus (O/A), and stratum pyramidale (PYR; Fig. 1b, c; Supplementary Table 1). Within this laminar projection pattern, MnR axonal varicosities were frequently observed in putative apposition with VIP-IN somata and dendrites (Fig. 1d; Supplementary Fig. 1e, f). These appositions were detected onto multiple VIP-IN subtypes, including CR+, CCK+ and CR–/CCK–VIP-INs (Fig. 1e, f). At the population level, CR+ VIP-INs accounted for the largest proportion of MnR-contacted VIP-INs, reflecting their greater abundance in the sampled dCA1 VIP-IN population (Fig. 1f, bottom). However, after accounting for the number of cells identified within each subtype, putative MnR appositions were detected in a high fraction in all VIP-IN subtypes (172/205 CR+ cells, 83.9%; 45/46 CCK+ cells, 97.8%; 35/38 CR−/CCK− cells, 92.1%; Fig. 1f, top), indicating broad targeting of multiple subtypes rather than exclusive per-cell selectivity for CR+ cells. Nevertheless, across layer and cellular compartments, the aggregate number of putative MnR appositions was highest onto CR+ VIP-INs, indicating that abundant VIP/CR-IS-3-like cells represent the dominant population-level target of MnR projections (Fig. 1g; Supplementary Table 1).

To determine whether this anatomical organization corresponds to functional connectivity, we performed optogenetic circuit mapping of MnR→VIP-IN inputs using whole-cell recordings from visually identified dCA1 VIP-INs. Post hoc immunostaining and morphological reconstruction identified three VIP-IN subtypes: VIP/CR-IS-3 cells, characterized by small somata and axons projecting to oriens/alveus (O/A; Fig. 1h); VIP/CCK-BCs, which displayed larger somata and axonal arbors concentrated in PYR (Fig. 1i); and CR/CCK-negative VIP-INs with somata located in O/A, hereafter referred to as VIP-OA cells, a heterogeneous group including, but not limited to, subiculum-projecting VIP/M2R-LRP cells (Tyan et al., 2014; Francavilla et al., 2018; Luo et al., 2019; Fig. 1j). Wide-field optical stimulation of MnR axons evoked short-latency excitatory postsynaptic currents (EPSCs) in all three VIP-IN subtypes, with similar connection probabilities (Fig. 1k, l; Supplementary Table 1). Among connected cells, EPSC amplitudes were largest and response latencies shortest in VIP/CCK-BCs (Fig. 1m, n; Supplementary Table 1), consistent with the prominent perisomatic appositions observed onto this subtype (Fig. 1g; Supplementary Fig. 1f). Thus, MnR projections provide functional excitatory input to multiple dCA1 VIP-IN subtypes, while engaging these populations with distinct response strength and timing.

### Glutamatergic and serotonergic mechanisms regulate MnR engagement of VIP-INs

We next asked which transmitter-defined MnR pathways may regulate dCA1 VIP-INs. To separately examine glutamatergic and serotonergic MnR projections, we expressed Cre-dependent fluorescent reporter in the MnR of VGLUT3-Cre and SERT-Cre mice (Fig. 2a, b). Both VGLUT3+ and SERT+ MnR projections formed putative appositions with CR+ and CCK+ VIP-INs (Fig. 2a–c) as well as CR–/CCK–VIP-INs (data not shown), indicating that both transmitter-defined MnR pathways are anatomically positioned to influence multiple VIP-IN subtypes. The number of VGLUT3+ and SERT+ appositions per cell was comparable between CR+ and CCK+ VIP-INs; however, VGLUT3+ terminals formed more numerous appositions onto CR+ VIP-INs than did SERT+ terminals (Fig. 2c; Supplementary Table 1). Thus, MnR input to VIP-INs includes both glutamatergic and serotonergic components, with VGLUT3+ projections contributing more prominently to the population-level innervation of CR+ VIP-INs.

**Figure 2.**
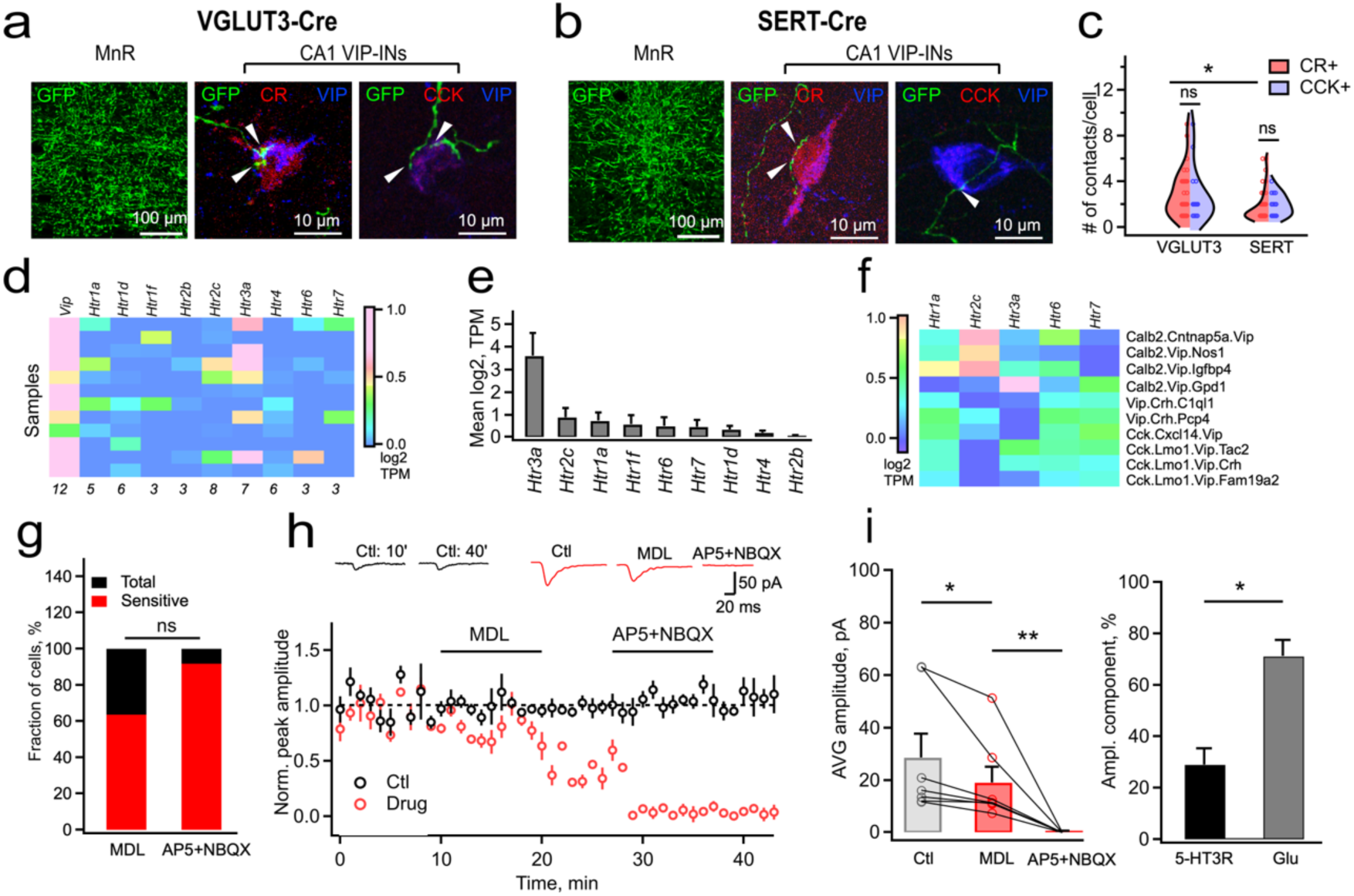
Glutamatergic and serotonergic MnR pathways excite CA1 VIP interneurons. **a,b,** Representative Cre-dependent labeling of VGLUT3-defined (**a**) and SERT-defined (**b**) MnR neurons and putative axonal appositions onto CA1 CR+ and CCK+ VIP-INs. Arrows indicate putative appositions. **c,** Number of VGLUT3+ and SERT+ putative appositions onto CR+ and CCK+ VIP-INs (VGLUT3, 86 CR+ and 38 CCK+ cells from 2 mice; SERT, 63 CR+ and 26 CCK+ cells from 3 mice; see Supplementary Table 1 for full statistical details). **d,e,** Single-cell Patch-seq expression of *Vip* and serotonin-receptor genes in 12 CA1 VIP-INs (**d**) and corresponding mean transcript abundance (**e**). **f,** Serotonin-receptor gene expression across VIP-IN subtypes in the Harris et al. (2018) CA1 transcriptomic dataset. **g,** Fraction of recorded VIP-INs showing an MDL-72222- and AP5+NBQX-sensitive MnR components. **h,** Representative MnR-evoked EPSCs and time courses during control recording, MDL-72222 application, and AP5 plus NBQX. **i,** Mean EPSC amplitudes across pharmacological conditions (left) and estimated glutamatergic and 5-HT3aR-sensitive response components (right; p = 0.0313, Wilcoxon rank test, n = 7 cells from 5 mice, 2 male/3 female). Full statistical details are provided in Supplementary Table 1. \**p < 0.05, **p < 0.01;* ns, not significant.

Because VIP-INs belong to a broader 5-HT3a receptor-expressing interneuron class and may also express metabotropic 5-HT receptors (Lee et al., 2013; Paul et al., 2017; Gouwens et al., 2020; Pronneke et al., 2020), we next examined serotonin receptor expression in CA1 VIP-INs. Patch-seq analysis of recorded VIP-INs revealed variable *Htr* gene expression across cells, with *Htr3a* detected at the highest expression levels, followed by *Htr2c* (Fig. 2d, e; Supplementary Fig. 2a–c). A similar receptor-expression profile was observed in VIP-INs from the Allen Brain Map mouse whole cortex and hippocampus SMART-seq dataset (Supplementary Fig. 2d). In addition, analysis of an independent hippocampal CA1 transcriptomic dataset further confirmed expression of major *Htr* genes across VIP-IN subtypes, including *Calb2*- and *Cck*-expressing populations (Fig. 2f; Harris et al., 2018). Together, these data indicate that both ionotropic 5-HT3a and metabotropic 5-HT2c receptors are positioned to regulate CA1 VIP-IN activity.

To test whether serotonin contributes directly to MnR-evoked synaptic responses, we examined the sensitivity of light-evoked MnR→VIP-IN EPSCs to the 5-HT3aR antagonist MDL-72222. MDL reduced MnR-evoked EPSCs in a subset of VIP-INs (7/11 cells), including identified VIP/CR-IS-3 cells and VIP/CCK-BCs (Fig. 2g–i). The remaining response was abolished by AP5 and NBQX, which also completely blocked the response in MDL-insensitive cells, indicating that the dominant component of fast MnR→VIP-IN transmission is glutamatergic, with an additional 5-HT3a-mediated contribution in a subset of cells (Fig. 2h, i; Supplementary Table 1).

We then asked whether serotonin also regulates VIP-IN excitability through metabotropic 5-HT2c receptors. Bath application of 5-HT increased VIP-IN excitability, reducing the action potential (AP) threshold and afterhyperpolarization (AHP) amplitude and increasing firing during somatic current injection, without changing AP amplitude or half-width (Supplementary Fig. 3; Supplementary Table 1). These effects were abolished when 5-HT was co-applied with the selective 5-HT2cR antagonist RS-102221 (Supplementary Fig. 3c, f, g). Thus, MnR projections regulate CA1 VIP-INs through a dominant fast glutamatergic excitation, a 5-HT3aR-mediated synaptic component, and 5-HT2cR-dependent enhancement of VIP-IN intrinsic excitability.

### Novelty preferentially recruits speed-coupled VIP-INs

The anatomical and ex vivo mapping experiments above indicate that MnR projections are positioned to recruit multiple CA1 VIP-IN subtypes, with the dominant population-level contact burden falling onto abundant VIP/CR-IS-3-like cells. We next asked how this anatomical organization relates to VIP-IN recruitment during behavior. VIP-INs are strongly activated during exploration of novel environments (Tamboli et al., 2024; Neubrandt et al., 2025), but this population is heterogeneous, and distinct VIP-IN subtypes may contribute differently to novelty processing. To resolve VIP-IN activity at single-cell resolution, we imaged dCA1 VIP-INs in freely moving Vip-IRES-Cre mice injected with pGP-AAV-CAG-Flex-jGCaMP7b-WPRE and implanted with a GRIN lens above dCA1 (Fig. 3a–d; Supplementary Fig. 4a, b).

**Figure 3.**
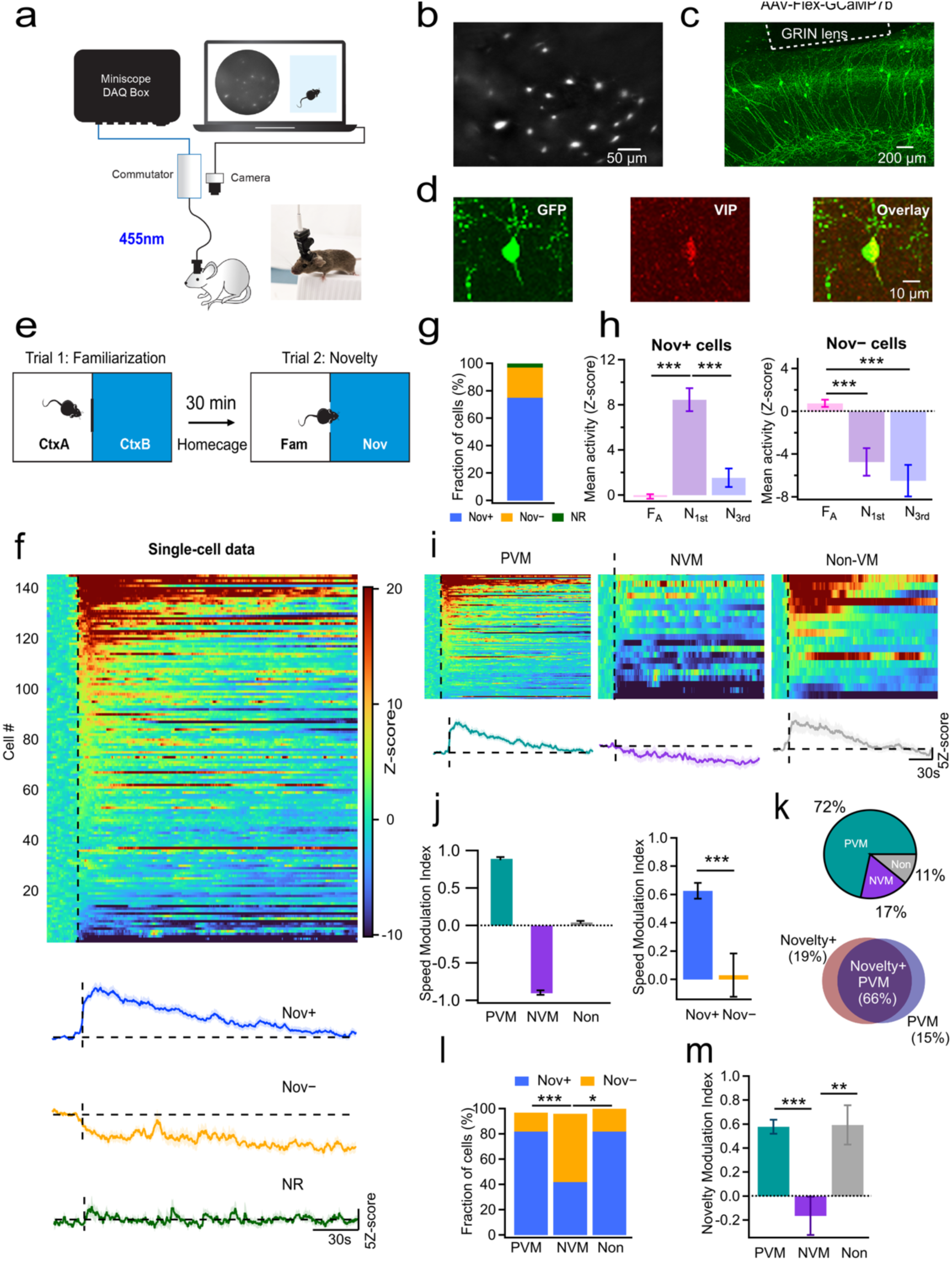
Novelty preferentially recruits positively velocity-modulated VIP interneurons. **a,** In vivo calcium-imaging configuration using the nVoke miniscope. **b,** Representative field of view showing GCaMP7b-expressing dCA1 VIP-INs. **c,** GCaMP7b expression and GRIN-lens placement above dCA1. **d,** GCaMP7b, VIP immunolabeling, and overlay. **e,** Familiar-to-novel context-transition paradigm. **f,** VIP-IN activity aligned to entry into the novel context (dashed line; n = 146 cells from 11 mice), with mean traces for novelty-activated (Nov+), novelty-suppressed (Nov−), and non-responsive (NR) cells. **g,** Fraction of cells in each response class. **h,** Activity profiles during the last minute in familiar context A (F_A_), the first minute in the novel context (N_1st_), and the third minute in the novel context (N_3rd_) (Nov+: F(2,216) = 67.938, p = 5.67 x 10^-20^; rmANOVA, n = 109 cells from 11 mice, 4 male/ 7 female; Nov–: χ^2^(2) = 41.312, p = 1.069 x 10^-9^; n = 32 cells from 8 mice, 2 male/ 6 female; see Supplementary Table 1 for full statistical details). **i,** Activity heat maps and mean traces of positively velocity-modulated (PVM; n = 105), negatively velocity-modulated (NVM; n = 25), and non-velocity-modulated (non-VM; n = 16) cells. **j,** Speed-modulation indices across velocity classes and between Nov+ and Nov− cells (p = 6.68 x 10^-5^; Wilcoxon rank test). **k,** Distribution of velocity classes and overlap between Nov+ and PVM classifications; percentages refer to the total number of cells. **l,** Fraction of Nov+ and Nov− cells within each velocity class (PVM: 105 cells, NVM: 25 cells, Non-VM: 16 cells; 11 mice, 4 male/ 7 female; see Supplementary Table 1 for full statistical details). **m,** Novelty-modulation index across PVM, NVM, and non-VM cells (H(2) = 18.4726, p = 9.74 x 10^-5^; Kruskal-Wallis test). Data are shown as mean ± s.e.m. where indicated. Full statistical details are provided in Supplementary Table 1. \*\**p < 0.01, ***p < 0.0001*.

Mice were tested in a contextual novelty-transition paradigm consisting of two distinct environmental layouts, Context A and Context B, separated by a wall with a hidden door (Fig. 3e). In the first trial, mice explored Context A for 5 min to establish familiarity. In the second trial, mice were re-introduced into the now-familiar Context A for 5 min, after which the door was opened to allow self-initiated entry into the novel Context B (Fig. 3e). Entry into the novel context evoked an immediate increase in activity in most VIP-INs (75%; n = 146 cells/11 mice; Fig. 3f, g), which gradually returned toward baseline as animals explored and familiarized with Context B. Novelty responses, however, were not uniform: a subset of VIP-INs was suppressed in the novel environment, whereas other cells showed little or no modulation (Fig. 3f). Based on a novelty modulation index, VIP-INs were therefore classified as novelty-excited (Nov+), novelty-inhibited (Nov–), or non-responsive (NR) cells (Fig. 3f, g). Specifically, Nov+ VIP-INs showed significantly higher activity during the first minute in the novel context than during familiar exploration and significantly reduced their activity over 3-min novel-context exploration (Fig. 3h, left; Supplementary Table 1). In contrast, Nov–VIP-INs were more active in the familiar context and remained suppressed for the entire duration of novel-context exploration (Fig. 3h, right; Supplementary Table 1).

Because CA1 VIP-IN subtypes differ in their relationship to locomotion, we next asked whether novelty-defined VIP-IN ensembles map onto previously described velocity-modulated populations. In head-restricted mice running on a treadmill, VIP-INs have been subdivided into cells positively modulated by running velocity (PVM) and negatively modulated by velocity (NVM) (Turi et al., 2019). These functional classes are associated with distinct VIP-IN subtypes, with PVM cells enriched among VIP/CR-IS-3-like cells and NVM cells including VIP/CCK-BCs and VIP-M2R/LRP-related populations (Francavilla et al., 2018; Luo et al., 2020; Geiller et al., 2020; Dudok et al., 2021). We confirmed this functional distinction on freely moving mice by correlating single-cell VIP-IN activity with animal running speed and calculating a speed modulation index in familiar context A (Fig. 3i, j; Supplementary Fig. 4c–e). Notably, cells identified as PVM in the familiar environment were enriched among Nov+ VIP-INs in the novel context, whereas NVM cells were enriched among Nov–VIP-INs (Fig. 3j–l; Supplementary Table 1). Non-velocity-modulated (Non-VM) VIP-INs showed a mixed profile with a dominant recruitment in novel environment (Fig. 3i, right; 3l, m; Supplementary Table 1). Thus, the VIP-IN ensemble recruited during novel-context exploration is biased toward the speed-upregulated, CR+/IS-3-like disinhibitory population. Together, these results indicate that novelty does not recruit CA1 VIP-INs uniformly. Instead, novel-context exploration preferentially engages a speed-upregulated VIP-IN ensemble, while suppressing a smaller group of speed-negatively modulated VIP-INs.

Because novel context exploration is accompanied by increase in running speed, we next tested whether the Nov+ response could be explained simply by increased locomotion. We compared VIP-IN activity in familiar and novel contexts at matched running velocities (Supplementary Fig. 5). Nov+ cells remained significantly more active in the novel context than in the familiar context at comparable speeds, indicating that their recruitment reflects novelty-related modulation beyond locomotor drive alone (Supplementary Fig. 5b, c). Thus, novelty preferentially recruits a speed-upregulated VIP/CR-IS-3-like VIP-IN population through a signal that is not reducible to running speed.

### MnR input supports robust recruitment of PVM VIP-INs during novel experience

The preceding analyses showed that novel-context exploration preferentially recruits a Nov+/PVM VIP-IN ensemble enriched for VIP/CR-IS-3-like cells, the population that receives the dominant aggregate MnR apposition burden with a prominent glutamatergic component (Fig. 2c). We next asked whether MnR input can support robust activation of this VIP-IN ensemble during novel experience. To this end, we combined cellular-resolution calcium imaging of dCA1 VIP-INs with simultaneous optogenetic inhibition of MnR→dCA1 projections in freely moving mice (Fig. 4). Vip-IRES-Cre mice were injected in dCA1 with pGP-AAV-CAG-Flex-jGCaMP7b-WPRE to enable imaging of VIP-INs, and in the MnR with a combination of AAV2-PHPeB-CaMKII-Cre-tdTomato and Cre-dependent Jaws-GFP to inhibit MnR→dCA1 projections (Fig. 4a, b; Supplementary Fig. 6a–d). Consistent with previous reports that CaMKII is expressed by both VGLUT3+ and serotonergic MnR neurons (Hioki et al., 2010; Ren et al., 2018), this strategy targeted both transmitter-defined MnR populations, with a bias toward VGLUT3+ cells (Supplementary Fig. 6c, d). This approach therefore enabled pathway-level suppression of MnR→CA1 inputs rather than transmitter-specific manipulation.

**Figure 4.**
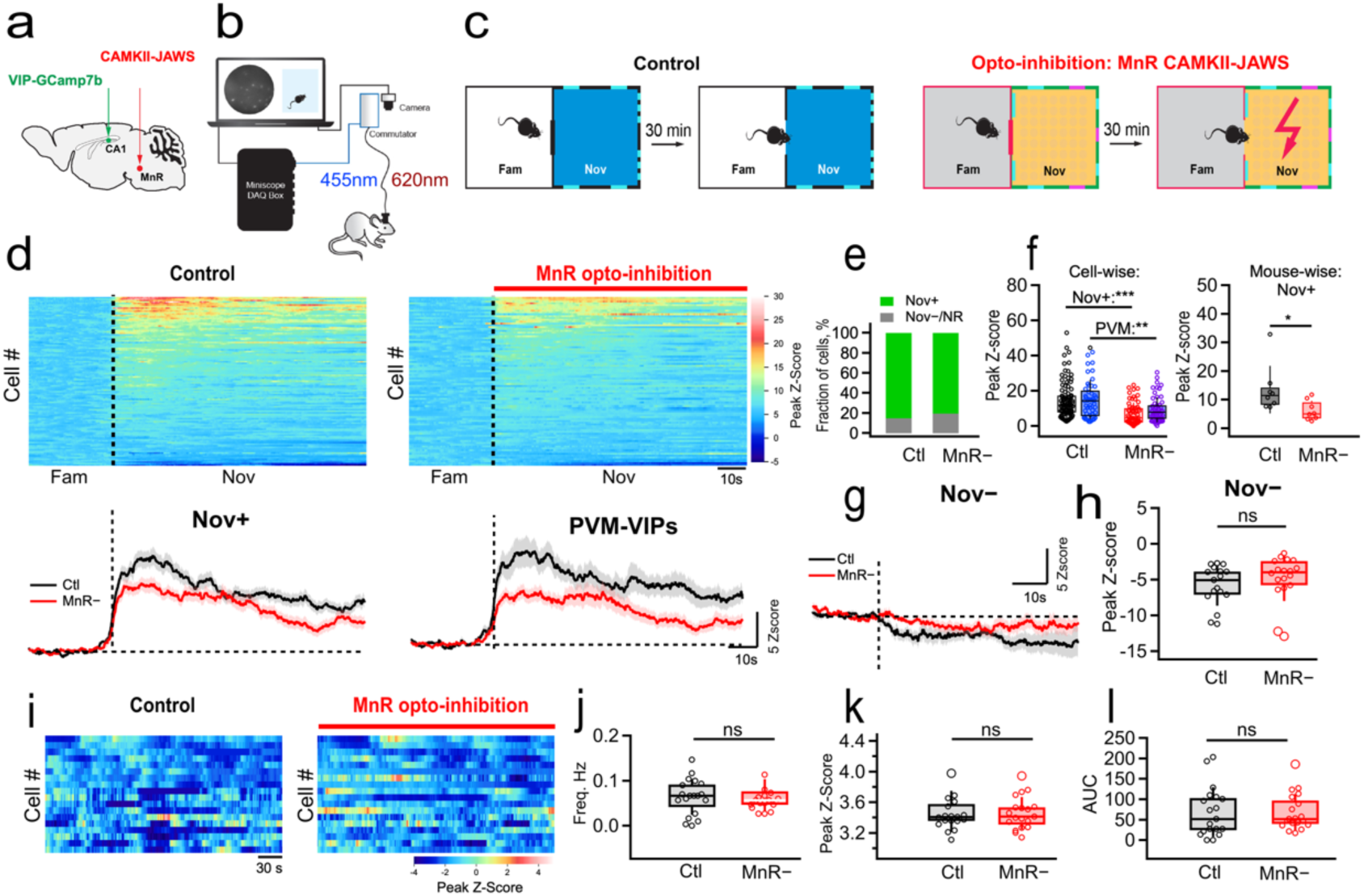
MnR→dCA1 projections support VIP-IN recruitment during novel-context exploration. **a,** Viral strategy for expression of GCaMP7b in dCA1 VIP-INs and Jaws in CaMKII-expressing MnR neurons. **b,** Experimental configuration for simultaneous single-cell calcium imaging and optogenetic inhibition of MnR→dCA1 projections in freely moving mice. **c,** Familiar-to-novel transition under control conditions and during MnR→dCA1 opto-inhibition in the novel compartment. **d,** VIP-IN activity aligned to novel-context entry under control and opto-inhibition conditions. Mean traces show Nov+ cells and PVM cells (n = 97 cells from 8 mice, 4 male/4 female). Dashed lines indicate novel-context entry. **e,** Fraction of cells corresponding to Nov+ vs. Nov–/non-responsive under each condition (p = 0.4561, two-sided Fisher exact test, n = 114 cells under control and 92 cells under MnR→dCA1 opto-inhibition, 8 mice, 4 male/4 female). **f,** Peak novelty-evoked activity of Nov+ cells (left, cell-wise p = 8.59 × 10⁻⁹, n = 97 cells from 8 mice; right, mouse-wise p = 0.0156, n = 8 mice, 4 male/ 4 female; Wilcoxon rank test), and of PVM cells (left, cell-wise p = 0.0036, n = 54 cells from 8 mice; Wilcoxon rank test) was reduced during MnR→dCA1 inhibition. **g,h,** Mean activity and peak responses of Nov− cells under control and MnR opto-inhibition conditions (p = 0.9283, Wilcoxon rank test, n = 35 cells, 4 mice, 2 male/2 female). **i,** VIP-IN activity during familiar open-field exploration under control and MnR opto-inhibition conditions. **j–l,** Calcium-transient frequency (**j**), peak amplitude (**k**), and area under the curve (**l**) (n = 18 cells from 3 mice, 2 male/1 female). Box plots show median and interquartile range; points represent cells. Full statistical details are provided in Supplementary Table 1. *\*\*\*p < 0.001*; ns, not significant.

Histological analysis further showed only sparse VIP-tdTomato+ neurons within the MnR and no detectable VIP+ projections to CA1 in Vip-IRES-Cre mice, excluding detectable contamination of CA1 terminal targeting by VIP-expressing MnR pathways (Supplementary Fig. 6e, f). In acute slices, activation of Jaws in MnR axons reduced VIP-IN Ca^2+^-transients evoked by electrical stimulation in SLM, confirming that the optogenetic protocol suppresses a MnR-sensitive component of synaptically driven VIP-IN recruitment (Supplementary Fig. 6g, h).

To test the contribution of MnR input to novelty-related VIP-IN recruitment, mice were exposed to a contextual novelty-transition paradigm using arenas with identical geometry but distinct sensory cues (Fig. 4c). Under control conditions, most VIP-INs showed a rapid increase in activity upon entry into the novel environment, consistent with the Nov+/PVM population identified above (Fig. 4d, top left). Based on the novelty modulation index, 97 out of 114 cells were classified as Nov+ under control conditions, whereas 74 out of 92 cells were classified as Nov+ during MnR→dCA1 opto-inhibition. Nevertheless, MnR→dCA1 opto-inhibition did not significantly alter the fraction of Nov+ cells (Fig. 4e; Supplementary Table 1), but markedly reduced the magnitude of the Nov+ response (Fig. 4d, bottom left; 4f). A similar reduction was observed in PVM VIP-INs, consistent with the strong overlap between Nov+ and PVM VIP-IN identity (Fig. 4d, f; Supplementary Table 1). This effect was selective for the Nov+ ensemble, as the activity profile of Nov− cells was not significantly altered (Fig. 4g, h; Supplementary Table 1). Importantly, inhibiting MnR→dCA1 projections during baseline exploration of a familiar open field did not alter VIP-IN Ca^2+^ transient frequency, amplitude, or area under the curve (Fig. 4i–l; Supplementary Table 1). Together, these results indicate that MnR input does not support the initial classification of VIP-INs as novelty responsive, nor does it tonically control VIP-IN activity during baseline exploration. Instead, MnR amplifies the recruitment of the Nov+/PVM VIP-IN ensemble during novel-context exploration.

To examine how MnR-dependent VIP-IN recruitment could influence downstream CA1 dynamics during novelty, we adapted a circuit model (Turi et al., 2019) to incorporate a familiar-to-novel transition during virtual mouse exploration in a two-dimensional space, novelty-related EC/CA3 and MnR inputs to multiple VIP-IN populations, and their downstream interactions with OLM interneurons, parvalbumin-expressing basket cells (PV-BCs), and pyramidal cells (Fig. 5a). In control simulations, the familiar-to-novel transition recruited VIP/CR-IS-3 and, to a lesser extent, VIP/CCK-BC and pyramidal cell activity while reducing OLM, PV-BC, and VIP-NVM activity, consistent with a shift toward circuit disinhibition during novelty (Fig. 5b, c). Global MnR removal dampened this transition, reducing VIP/CR-IS-3 recruitment, preventing OLM suppression, and decreasing place-cell density (Fig. 5c, left; d; Supplementary Table 1). Notably, selective removal of MnR input to VIP/CR-IS-3 largely reproduced the effects of global MnR removal on VIP/CR-IS-3 activity, OLM activity, and place-cell density (Fig. 5c, left; d; Supplementary Table 1), whereas removal of MnR input to VIP/CCK-BC and VIP-NVM populations enhanced PV-BC activity (Fig. 5c, bottom middle). In addition, VIP/CCK-BC and VIP-NVM activity increased when MnR input to VIP/CR-IS-3 was selectively removed (Fig. 5c, top middle and right; Supplementary Table 1), consistent with a negative influence of the VIP/CR-IS-3 pathway onto VIP/CCK-BCs and VIP-NVMs in the model. These simulations support a model in which MnR input facilitates novelty-related CA1 output primarily through recruitment of the VIP/CR-IS-3→OLM disinhibitory pathway, with VIP/CCK-BC and VIP-NVM populations regulated by a balance between direct MnR excitation and indirect inhibition through MnR-driven VIP/CR-IS-3 recruitment.

**Figure 5.**
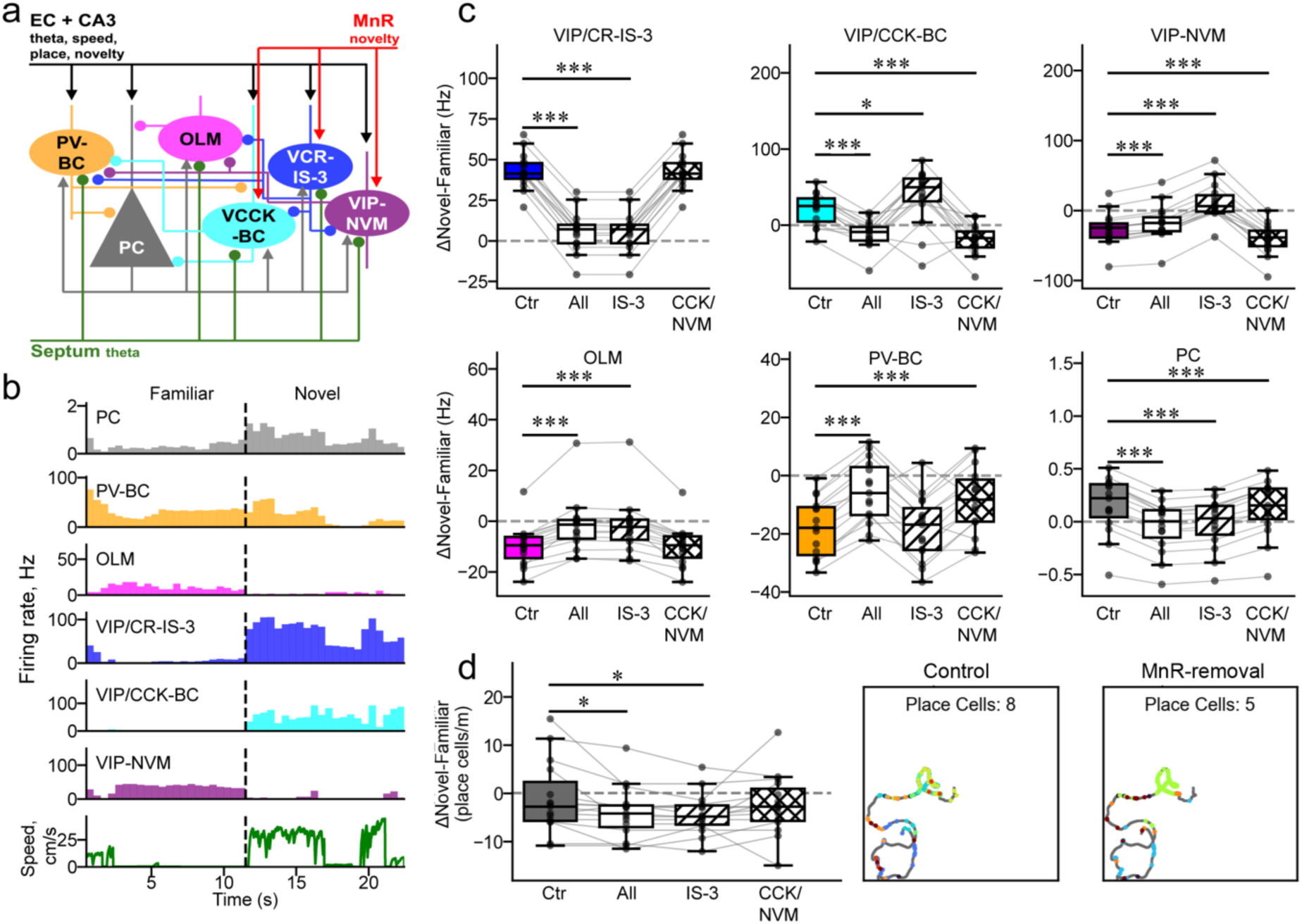
Simulated MnR input modulates VIP disinhibitory microcircuits and CA1 place-cell output during spatial novelty. **a,** Schematic of the hippocampal CA1 microcircuit model incorporating pyramidal cells (PCs), parvalbumin-expressing basket cells (PV-BC), oriens-lacunosum moleculare cells (OLM), VIP/CCK basket cells (VCCK-BC), VIP/CR-IS-3 positively velocity-modulated cells (VCR-IS-3), and VIP negatively velocity-modulated cells (VIP-NVM). Axo-axonic and bistratified cells were included in the model but are not shown. The network receives entorhinal cortex (EC), CA3, septal theta, and novelty-modulated MnR inputs. **b,** Example population firing-rate histograms (500-ms bins) and running-speed profile during transition from familiar to novel context in a 100 × 100-cm arena under control conditions. The dashed line indicates novel-context entry. **c,** Familiar-to-novel firing-rate changes under control conditions (Ctr), global MnR-input removal (All), selective removal from VIP/CR-IS-3 cells (IS-3), and combined removal from VIP/CCK-BC and VIP-NVM populations (CCK/NVM). Full statistical details are provided in Supplementary Table 1. **d,** Change in CA1 place-cell density across the same conditions. Insets show representative trajectories and place-cell spikes under control and global MnR-input removal. Box plots show median and interquartile range; whiskers extend to 1.5 × IQR. Points represent simulation runs, and gray lines connect runs generated with matched random seeds (n = 15 virtual trials per condition). *\*p < 0.05, ***p < 0.001*.

Because MnR activity has also been implicated in stress-related behaviors (Balazsfi et al., 2017; Bocchio et al., 2016; Szonyi et al., 2019), we also assessed VIP-IN responses during the tail-suspension test. Tail suspension robustly activated VIP-INs under control conditions, but this activation was unaffected by MnR→dCA1 opto-inhibition (Supplementary Fig. 7), indicating that MnR-dependent VIP-IN recruitment is behavioral-context specific. Together, these results show that MnR projections support robust recruitment of the novelty-activated CA1 VIP-IN ensemble and suggest that this effect preferentially engages a VIP-PVM→OLM disinhibitory motif, while being dispensable for baseline exploratory and stress-related VIP-IN activation.

### MnR→dCA1 projections constrain novelty-guided exploratory sampling and support spatial location memory

The imaging and modeling results above indicate that MnR→dCA1 input preferentially amplifies recruitment of the Nov+/PVM VIP/CR-IS-3 ensemble during novel experience and may thereby influence CA1 output during novelty. We next asked whether this circuit modulation is associated with a behavioral consequence during exploration of a novel environment. We therefore analyzed exploratory behavior during the contextual novelty-transition paradigm under control conditions and during optogenetic inhibition of MnR→dCA1 projections restricted to the initial phase of novel-context exploration (Fig. 6a).

**Figure 6.**
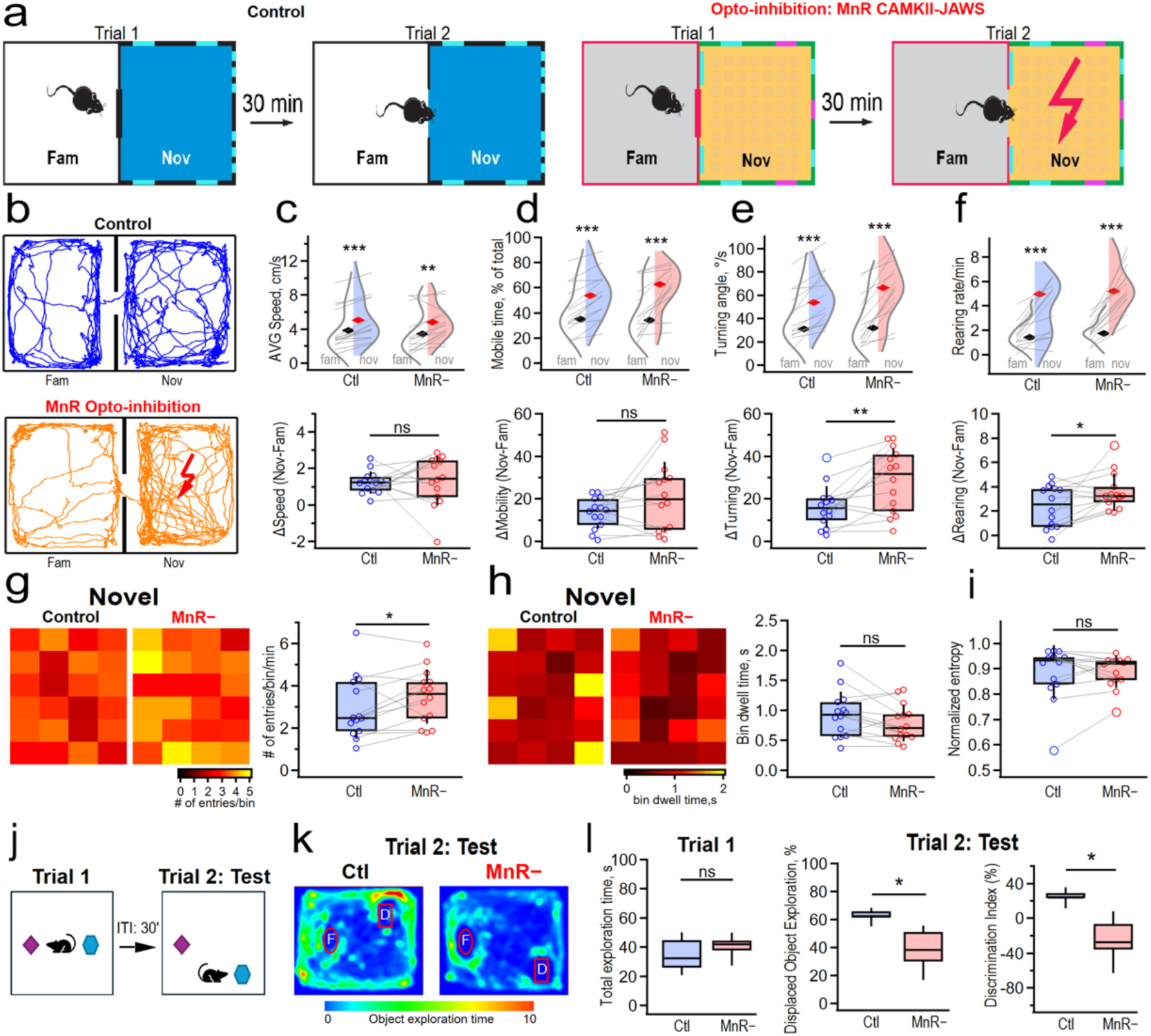
MnR→dCA1 projections regulate novelty-guided exploratory sampling and support object-location memory. **a,** Familiar-to-novel transition under control conditions and during optogenetic inhibition of MnR→dCA1 projections. During opto-inhibition sessions, 620-nm illumination was delivered for 90 s beginning upon entry into the novel compartment. **b,** Representative locomotor trajectories. **c–f,** Average speed (**c**), mobile time (**d**), absolute turning angle (**e**), and rearing rate (**f**) during familiar and novel exploration, with novelty-associated changes shown below. MnR→dCA1 photo-inhibition did not detectably alter the novelty-associated change in speed or mobile time but increased the change in turning angle (**e**, p = 0.0031, Wilcoxon signed-rank test) and rearing rate (**f**, p = 0.0419, Wilcoxon signed-rank test) (n = 14 mice, 6 male/8 female). See Supplementary Table 1 for full statistical details. **g,** Entries per 6-cm spatial bin during novel exploration (p = 0.0494, Wilcoxon signed-rank test; n = 14 mice, 6 male/8 female). **h,** Bin dwell time (p = 0.104, Wilcoxon signed-rank test). **i,** Normalized spatial entropy across 24 bins (p = 0.9153, Wilcoxon signed-rank test). **j,** Object-location task. MnR→dCA1 projections were inhibited during the sample trial; memory was tested after 30 min. **k,** Representative test-trial occupancy maps. F, familiar-location object; D, displaced object. **l,** Total sample-phase exploration (left), test-phase exploration of the displaced object (middle; p = 0.03125, Wilcoxon signed-rank test), and discrimination index (right; p = 0.0156, Wilcoxon signed-rank test; n = 6 mice, 3 male/3 female). Violin and box plots show individual mice; gray lines connect paired observations. Box plots show median and interquartile range. Full statistical details are provided in Supplementary Table 1. *p < 0.05, **p < 0.01, ***p < 0.001; ns, not significant.

Transition from the familiar to the novel context induced a robust exploratory response in both control and MnR opto-inhibition sessions (Fig. 6b). In both conditions, mice increased average speed, percentage of mobile time, turning angle, and rearing upon entry into the novel context (Fig. 6c–f), indicating that MnR→dCA1 photo-inhibition did not prevent animals from detecting or engaging with the novel environment. However, the structure of novelty-guided exploration was altered during MnR→dCA1 opto-inhibition. Although the novelty-induced increase in average speed and mobile time did not differ significantly between control and opto-inhibition sessions (Fig. 6c, d; bottom), MnR opto-inhibition significantly enhanced the novelty-induced increase in turning angle and rearing (Fig. 6e, f; bottom; Supplementary Table 1). Thus, MnR→dCA1 opto-inhibition did not reduce exploratory drive; instead, it enhanced specific components of novelty-guided spatial reorientation and active sampling.

To determine whether this altered exploratory pattern was reflected in spatial sampling of the novel environment, we divided the novel compartment into 6-cm spatial bins and quantified bin entries, dwell time, and occupancy entropy during novel-context exploration. MnR→dCA1 opto-inhibition significantly increased the number of entries per spatial bin (Fig. 6g), indicating more frequent transitions across the spatial layout. In contrast, the mean dwell time per bin visit was not significantly altered (Fig. 6h), and normalized spatial entropy calculated from the dwell-time distribution across bins was unchanged (Fig. 6i; Supplementary Table 1). Thus, MnR→dCA1 opto-inhibition did not alter overall novelty-driven locomotor engagement, but selectively increased spatial reorientation, rearing, and spatial sampling transitions, suggesting that MnR→dCA1 input normally helps optimize novelty-guided exploration.

Importantly, these behavioral effects were novelty specific. During baseline open-field exploration, MnR→dCA1 opto-inhibition did not alter average speed, mobile time, immobility events, turning angle, rearing, or thigmotaxis (Supplementary Fig. 8a). The open field tested tonic MnR→dCA1 contribution during familiar baseline exploration, whereas the familiar-to-novel transition tested MnR contribution during a self-initiated transition from a known context into a new spatial context. To control for the transition itself, we added a familiar-to-familiar transition using the same door-opening and optogenetic timing. MnR→dCA1 inhibition in this paradigm did not alter speed, mobile time, turning angle, or rearing (Supplementary Fig. 8b, c). Together, these data indicate that the behavioral changes observed during the familiar-to-novel transition cannot be explained by nonspecific effects of light delivery, altered baseline locomotion, anxiety-like spatial preference, or movement between compartments. Instead, they reveal a novelty-dependent role for MnR→dCA1 projections in shaping exploratory strategy. Finally, we asked whether MnR→dCA1 activity during encoding contributes to subsequent spatial-location memory, a process that depends on VIP-IN activity (Tamboli et al., 2024). In the object-location memory task, MnR→dCA1 projections were optogenetically inhibited during the sample trial, when mice explored two similar objects, and memory was assessed after a 30-min delay by relocating one object (Fig. 6j, k; Supplementary Table 1). Opto-inhibition of MnR→dCA1 projections did not affect total object exploration during the sample trial (Fig. 6l, left), indicating that object sampling during encoding was maintained. However, opto-inhibited mice showed reduced exploration of the displaced object during the test trial (Fig. 6l, middle), and a lower object-location discrimination index (Fig. 6l, right; Supplementary Table 1). Thus, MnR→dCA1 input during encoding supports the formation of spatial-location memory.

Taken together, these behavioral data establish a functional consequence of MnR→dCA1 recruitment during novel experience. MnR→dCA1 projection inhibition did not detectably alter general locomotion, transition between compartments, or the initiation of novelty exploration. Instead, MnR inputs constrain the structure of novelty-guided sampling and support the encoding of spatial-location memory.

## Discussion

This study identifies a brainstem–hippocampal circuit motif that regulates dCA1 VIP-IN dynamics and spatial exploration during novel experience. We show that MnR projections regulate multiple dCA1 VIP-IN subtypes through glutamatergic and serotonergic signaling, with the dominant population-level anatomical burden falling onto abundant CR+/IS-3-like VIP-INs. During behavior, novel-context exploration preferentially recruits a Nov+/PVM VIP-IN ensemble enriched for this CR+/IS-3-like population. MnR→dCA1 opto-inhibition does not abolish novelty responsiveness, but selectively reduces the magnitude of Nov+/PVM VIP-IN recruitment, alters novelty-guided spatial sampling, and impairs object-location memory when applied during encoding. Thus, MnR input does not simply excite hippocampal VIP-INs; it amplifies a specific disinhibitory state that supports organized exploration and spatial encoding.

A central conclusion is that MnR projections are organized to influence several VIP-dependent inhibitory motifs in parallel. Consistent with previous anatomical work showing dense raphe innervation of hippocampal CA1 (Freund et al., 1990; Papp et al., 1999; Varga et al., 2009; Szonyi et al., 2016; Fortin-Houde et al., 2023), we found that MnR axons contact VIP-IN somata and dendrites across CA1 layers. At the population level, CR+ VIP-INs accounted for the largest share of MnR-contacted cells and the greatest aggregate number of putative appositions. Functionally, however, MnR-evoked EPSCs were detected across VIP-IN subtypes, with the largest and fastest responses observed in VIP/CCK-BCs. MnR input is therefore not restricted to one VIP-IN subtype, but distributed across multiple inhibitory and disinhibitory populations, with circuit impact likely determined by connectivity motifs and specific behavioral state during which these motifs are activated.

MnR modulation of VIP-INs is also transmitter-complex, with both VGLUT3+ and serotonergic projections contacting hippocampal VIP-INs, consistent with the mixed transmitter organization of the MnR and with the hippocampus being an established output target of both MnR VGLUT3+ and serotonergic neurons (Jackson et al., 2009; Varga et al., 2009; Hioki et al., 2010; Sos et al., 2017; Ren et al., 2019; Szonyi et al., 2016; Fortin-Houde et al., 2023). Functionally, MnR→VIP-IN responses contained a dominant glutamatergic component and, in a subset of cells, a 5-HT3aR-sensitive synaptic component. In addition, serotonin increased VIP-IN excitability through a 5-HT2cR-sensitive mechanism, consistent with known serotonergic actions on persistent Na⁺ currents, M-type K⁺ currents, and SK channels (Harvey et al., 2006; Deemyad et al., 2011; Roepke et al., 2012). This combination of fast excitation and modulatory gain control is well suited for novelty processing: glutamate and 5-HT3aR signaling can rapidly recruit VIP-INs at the onset of salient experience, whereas 5-HT2cR-dependent modulation may prolong or stabilize VIP-IN responsiveness during ongoing exploration.

During novelty, this pathway preferentially amplifies a functionally defined VIP-PVM ensemble. VIP-INs can be subdivided into populations positively or negatively modulated by running velocity, with PVM cells enriched among VIP/CR-IS-3 cells and NVM cells including VIP/CCK-BCs and VIP-M2R/LRP-related populations (Francavilla et al., 2018; Turi et al., 2019; Luo et al., 2020; Geiller et al., 2020; Dudok et al., 2021). We found that Nov+ cells were enriched among PVM VIP-INs, whereas Nov− cells were enriched among NVM populations. Importantly, Nov+ VIP-INs remained more active in the novel context than in the familiar context at matched running speeds. Thus, novelty recruitment cannot be explained solely by enhanced locomotion. Novelty and movement converge on overlapping but separable VIP-IN ensembles, with MnR input preferentially strengthening the novelty-associated PVM ensemble response. This effect was absent during familiar open-field exploration, familiar-to-familiar transition and during tail-suspension-induced VIP-IN activation, indicating that MnR→dCA1 input does not act as a tonic regulator of VIP-IN activity across behavioral states, but mainly supports full-magnitude PVM VIP-IN recruitment during novel-context exploration.

The circuit model provides a mechanistic interpretation of this selectivity. In the model, the familiar-to-novel transition recruited VIP/CR-IS-3, VIP/CCK-BCs, and pyramidal cells, while strongly suppressing OLM activity and producing smaller changes in PV-BC and VIP-NVM activity. This pattern is consistent with a novelty-associated shift dominated by dendritic disinhibition, with additional modulation of perisomatic inhibitory balance through VIP/CCK-BC and PV-BC interactions. Global MnR removal dampened this novelty-transition effects and reduced place-cell density, while selective removal of MnR input to VIP/CR-IS-3 cells largely reproduced the effects of global MnR-removal on VIP/CR-IS-3 activity, OLM activity, and place-cell output. The model therefore places the MnR→VIP/CR-IS-3→OLM pathway at the core of MnR-dependent novelty processing in dCA1. At the same time, MnR engagement of VIP/CCK-BC and VIP-NVM populations provides additional circuit leverage through parallel inhibitory motifs, including regulation of perisomatic inhibition via the VIP/CCK-BC – PV-BC axis (Dudok et al., 2021) and modulation of dendritic inhibition and hippocampal-subicular coordination through VIP/M2R-LRP circuits (Francavilla et al., 2018). Importantly, this circuit balance may explain why VIP/CCK-BCs, despite receiving strong direct MnR excitation ex vivo, are not preferentially recruited as Nov+ cells in vivo: their activity during novelty likely reflects both direct MnR excitation and indirect regulation through the MnR-driven VIP/CR-IS-3 pathway.

This circuit logic predicts that perturbing MnR→dCA1 input should alter how novelty is explored, rather than simply whether novelty is detected. Consistent with this idea, MnR→dCA1 opto-inhibition did not prevent animals from engaging with novelty. Instead, mice explored the novel environment in a more effortful and fragmented manner, showing increased spatial reorientation, rearing, and sampling transitions without broad changes in locomotor engagement or occupancy structure. This phenotype was absent during baseline open-field exploration and during familiar-to-familiar transition using the same door-opening and light-delivery procedures, arguing against altered locomotion, anxiety-like behavior, light delivery, or transition itself as explanations. Thus, MnR→dCA1 input helps structure novelty-guided exploration, promoting efficient spatial sampling while preserving overall exploratory engagement.

These findings extend the known behavioral repertoire of MnR circuits. Although MnR activity has been linked prominently to anxiety- and stress-related behaviors (Bocchio et al., 2016; Balazsfi et al., 2017; Szőnyi et al., 2019; Pizzocarro et al., 2026), our dorsal CA1 manipulation revealed a spatially organized phenotype rather than a broad anxiety- or stress-like effect. This points to a hippocampal-axis-specific role for MnR output: ventral hippocampal projections may be more closely tied to affective regulation, whereas dorsal CA1 projections support exploration, spatial sampling, and memory encoding. Consistent with this view, both VGLUT3+/5-HT+ and VGLUT3+/5-HT–MnR neurons show elevated activity during dCA1 theta compared with non-theta states, while VGLUT3+/5-HT–population is strongly recruited by sensory stimulation (Domonkos et al., 2016), providing a potential mechanism by which raphe input could link sensory-driven exploratory states to hippocampal spatial computations.

This organization appears important for memory encoding. When MnR→dCA1 projections were inhibited during the sample phase of the object-location task, subsequent spatial-location memory was impaired. This result links MnR-dependent VIP-IN recruitment to a behavioral outcome requiring hippocampal encoding. A plausible mechanism is that MnR input gates dendritic disinhibition through VIP/CR-IS-3 recruitment at the moment when animals sample novel spatial configurations. Without this gain control, mice still explore, but exploration becomes less structured and less effective for encoding object location.

One limitation of our study is that the optogenetic strategy inhibited MnR→dCA1 input at the pathway level, impacting both glutamatergic and serotonergic components, with a bias toward VGLUT3+ neurons. This manipulation reflects the native mixed-transmitter architecture of the MnR pathway but does not isolate pure glutamatergic, serotonergic, or dual-transmitter projections. Dissecting these components in vivo will require complex intersectional and projection-specific strategies that simultaneously resolve MnR transmitter identity and VIP-IN activation, ideally at the level of defined subtypes.

Together, our findings place MnR→dCA1 projections at the interface between brainstem neuromodulatory state signals and hippocampal inhibitory microcircuits. MnR input engages CA1 VIP-INs through coordinated glutamatergic and serotonergic mechanisms, amplifies a novelty-recruited PVM VIP-IN ensemble, and supports exploratory structure and spatial-location memory. Rather than driving movement or novelty detection per se, MnR→dCA1 projections tune the inhibitory state of CA1 so that novel experience can be sampled and encoded effectively.

## Materials and Methods

### Mice

Experiments were performed using five mouse lines of either sex: VIP/enhanced green fluorescent protein (VIP-eGFP; Tyan et al., 2014) mice (BAC line with multiple gene copies; MMRRC strain #31009, STOCK Tg(Vip-EGFP) 37Gsat, University of California, Davis, CA), Vip-IRES-Cre mice, Vip-IRES-Cre-Ai9(tdTomato) mice, SERT-Cre mice (B6.FVB(Cg)-Tg(Slc6a4-cre)ET33Gsat/Mmucd (MMRRC, stock #031028-UCD), and VGLUT3-Cre mice, generously provided by Dr El Mestikawy (Centre de Recherche de l’Hôpital Douglas, Québec, Canada). The Vip-IRES-Cre-Ai9(tdTomato) mice were generated by crossing Vip-IRES-Cre mice (RRID:IMSR_JAX:010908, strain #010908, The Jackson Laboratory, Bar Harbor, ME, USA) with the Ai9 reporter line B6.Cg-Gt(ROSA)26Sortm9(CAG-tdTomato)Hze/J (RRID:IMSR_JAX:007909, stock #007909, The Jackson Laboratory, Bar Harbor, ME, USA). Vip-IRES-Cre, SERT-Cre, and VGLUT3-Cre mice were maintained heterozygotes on a C57BL6/J background. Mice were housed under standard conditions with ad libitum access to food and water and maintained on a 12 h light/dark cycle. Experiments were performed in accordance with the guidelines of the Animal Protection Committee of Université Laval and the Canadian Council on Animal Care under approved protocols 2019-148, 2022-1142, 2022-1160, and 2024-1865.

### Stereotaxic viral injections

Viral vectors were injected into the MnR region to label or manipulate MnR projections in hippocampal dCA1. For optogenetic mapping experiments, AAV5-hSyn-ChR2(H134R)-EYFP.WPRE.hGH (UNC Vector Core) was injected into the MnR of VIP-Cre-Ai9(tdTomato) mice. For transmitter-defined MnR projection labeling, AAVdj-EF1a-DIO-hChR2(E123T/T159C)-eYFP (6.5E12 GC/ml; Canadian Neurophotonics Platform Viral Vector Core Facility, RRID:SCR_016477) was injected into the MnR of SERT-Cre or VGLUT3-Cre mice. Mice were anesthetized with either ketamine-xylazine (100 mg/kg and 10 mg/kg, intraperitoneal) or isoflurane (3% for induction in oxygen, ∼0.8–1.5 L/min; 1.5–2% for maintenance in oxygen, ∼0.5 L/min) and placed in a stereotaxic frame (Kopf Instruments). In Vip-Cre-Ai9(tdTomato) mice, virus was injected into the MnR using the following coordinates relative to bregma: AP –4.24 mm, ML 0 mm, DV –4.5 mm. A total volume of 50–100 nl was delivered using a microprocessor-controlled nanoliter injector (Word Precision Instruments). In SERT-Cre and VGLUT3-Cre mice, 25–35 nl of virus was injected at 1 nl/s using a Nanoject III programmable nanoliter injector (Drummond Scientific) at AP –4.18 or –4.48, ML 0 mm, and DV –4.8. The injection capillary was inserted with a 10^◦^ lateral angle to avoid damaging the confluence of sagittal and transverse sinuses. After injection, the capillary was left in place for 5 min before slow withdrawal, and the scalp was sutured.

For in vivo calcium imaging, Vip-IRES-Cre mice were injected with pGP-AAV-CAG-Flex-jGCaMP7b-WPRE (Vigene Bioscience; Cat #: BS-12-CXBAAV9) in dCA1 using the following coordinates: AP –2.2 mm, ML –2.0 mm, DV –1.45 mm. For combined calcium imaging and optogenetic inhibition of MnR**→**CA1 projections, mice additionally received a MnR injection of a 1:1 mixture of AAV2-PHPeB-CaMKII-Cre-tdTomato (Viral Vector Core, Canadian Neurophotonics Platform) and pAAV5-CAG-Flex-rc [Jaws-KGC-GFP-ER2] (Addgene; Cat #: 84445). Two weeks after viral injection, mice were implanted with a GRIN lens (ProView™ Integrated Lens, Inscopix) above dCA1 for single-cell calcium imaging.

### Immunohistochemistry and anatomical analysis

For anatomical analysis of putative MnR contacts onto hippocampal dCA1 VIP-INs, Vip-IRES-Cre-Ai9(tdTomato) mice were injected in the MnR with AAV5-hSyn-ChR2(H134R)-EYFP and allowed to recover for 4–5 weeks. Mice were deeply anesthetized and perfused intracardially with sucrose-based artificial cerebrospinal fluid (ACSF), followed by 4% paraformaldehyde (PFA) and 20% picric acid in phosphate-buffered saline (PBS). Brains were removed and post-fixed overnight at 4 °C in 4% PFA/picric acid solution. The following day, brains were embedded in 4% agar and sectioned coronally at 40–70 µm using a vibratome (VT1000; Leica Microsystems or PELCO EasySlicer). Sections were stored in phosphate buffer containing 0.05–0.1% sodium azide.

Sections were permeabilized with 0.3% Triton X-100 in PBS and incubated for 1 h in blocking solution containing 20% normal serum. Sections were then incubated with primary antibodies for 24–48 h at 4°C, followed by incubation with fluorophore-conjugated secondary antibodies for 2–4 h at room temperature. Primary antibodies were: chicken anti-GFP (1:1000; Aves Labs Inc., Cat: GFP-1020), rabbit anti-5-HT (1:5000; Sigma, S-5545), guinea pig anti-VGLUT3 (1:250; EMD Millipore, Cat: AB5421-1), goat anti-calretinin (1:1000; Santa Cruz Biotechnology, Cat: sc-11644 or SWANT, Cat: CG1), rabbit anti-CCK (1:1000; Sigma, Cat: C2581), and guinea pig anti-VIP (1:1000; Synaptic Systems, Cat: 443 005). Secondary antibodies included donkey anti-chicken Alexa Fluor-488 (1:1000, Jackson Immunoresearch, Cat: 703-545-155), donkey anti-goat Dylight-650 (1:250; Thermo Scientific, Cat: SA5-10089), donkey anti-goat Alexa Fluor-546 (1:250; Thermo Scientific, Cat: A11056), donkey anti-rabbit Alexa Fluor-546 (1:250; Thermo Scientific, Cat: A10040), donkey anti-rabbit Alexa Fluor-647 (1:250; Invitrogen, Cat: A31573), and donkey anti-guinea pig Alexa Fluor-647 (1:250; Jackson Immunoresearch, Cat: 706-605-148).

Confocal images were acquired using a Leica TCS SP5 microscope equipped with 488-nm argon, 543-nm HeNe, or 633-nm HeNe lasers and 20x/0.8 NA or 63x/1.4 NA oil-immersion objectives. CA1 layers were manually delineated as stratum oriens/alveus (O/A), stratum pyramidale (PYR), stratum radiatum (RAD), and stratum lacunosum-moleculare (SLM), using NeuN immunostaining and the Allen Mouse Brain Atlas as anatomical guides. Quantification was restricted to dorsal-to-intermediate hippocampal sections to reduce septotemporal variability. Animals were included only when MnR viral labeling was consistent, defined as the presence of eYFP-positive axons in ≥80% of analyzed hippocampal sections. Axonal density was quantified within 123 × 123 µm regions of interest placed in the center of each CA1 layer and measured as mean eYFP fluorescence intensity (Fig. 1c). Putative MnR contacts onto VIP-INs were analyzed from high-resolution confocal z-stacks acquired with the 63× objective. Putative appositions were defined as close apposition between eYFP-positive MnR axonal varicosities and tdTomato-positive VIP-IN somata or dendrites, confirmed as overlap in orthogonal *x–y*, *x–z*, and *y–z* planes (Supplementary Fig. 1e). Dendritic appositions were counted only when the dendrite could be traced back to the soma. Somatic and dendritic appositions were quantified separately. The same quantification approach was applied to VIP-INs coexpressing CR or CCK.

For *post hoc* morphological identification of recorded VIP-INs, cells were filled with biocytin (0.6 mg/ml, Sigma) during whole-cell recordings. Slices were fixed overnight at 4°C in 4% PFA, permeabilized with 0.3% Triton X-100, and incubated with streptavidin-conjugated Alexa-488 (1:1000, Jackson Immunoresearch, Cat: 016-540-084) or Alexa-546 (1:1000, Invitrogen, Cat: S11225). Z-stacks were acquired with a 1-µm step size, and cells were reconstructed using Neurolucida.

### Patch-clamp recordings, optogenetic stimulation and two-photon calcium imaging ex vivo

Acute hippocampal slices were prepared from mice aged P79-P121. Mice were deeply anaesthetized using ketamine-xylazine and perfused intracardially with ice-cold high-sucrose ACSF as described previously (Francavilla et al., 2020; Michaud et al., 2024). Transverse hippocampal slices, 300 µm thick, were cut using a vibratome (Microm, Fisher Sci.) and transferred to a recording chamber continuously perfused with oxygenated ACSF containing (in mM): 124 NaCl, 2.5 KCl, 1.25 NaH_2_PO_4_, 26 NaHCO_3_, 2 MgSO_4_, 2 CaCl_2_, and 10 glucose (290-310 mOsm/L, pH 7.4). Recordings were performed at 30–32 °C. tdTomato-expressing VIP-INs were identified under epifluorescence using an upright Nikon Eclipse FN1 microscope equipped with a 40x/0.8NA objective. Voltage-clamp recordings were conducted using a cesium-based internal solution containing (in mM): 130 CsMeSO_4_, 2 CsCl, 10 diNa-phosphocreatine, 10 HEPES, 4 ATP-Tris, 0.4 GTP-Tris, 0.3% biocytin, 2 QX-314, 0.1 spermine, pH 7.2–7.3, 280–290 mOsm/L. Current-clamp recordings were conducted using a potassium (K^+^)-based internal solution containing (in mM): 130 K-gluconate, 2 KCl, 10 diNa-phosphocreatine, 10 HEPES, 4 ATP-Tris, 0.4 GTP-Tris, 0.3% biocytin, pH 7.2–7.3, 280–290 mOsm/L. Recordings were excluded if the access resistance exceeded 25 MΩ or if it changed by more than 15% during the experiment.

To assess monosynaptic inputs from MnR to dCA1 VIP-INs, voltage-clamp recordings were performed at a holding potential of –70 mV in the presence of 4-aminopyridine (4-AP; 1mM, Sigma, Cat #: 275875) and tetrodotoxin (1 µM, Alomone Labs, Cat #: T-550). ChR2-expressing terminals were activated with single blue light pulses delivered through the microscope objective using wide-field illumination (450–490 nm, 10 mW, 10 ms; David and Topolnik, 2017). Light stimuli were applied at 30-second intervals. Light-evoked EPSC peak amplitude was measured as the maximal inward current relative to the pre-stimulus baseline. Response latency was defined as the time from the end of the blue-light pulse to the onset of the light-evoked response, determined as the first deflection exceeding baseline noise and verified manually.

For *ex vivo* validation of Jaws-mediated inhibition of MnR projections, acute hippocampal slices were prepared from mice expressing GCaMP7b in CA1 VIP-INs and Jaws-eGFP in MnR neurons. VIP-IN Ca²⁺ transients were imaged using a Leica SP5 two-photon microscope equipped with a 25x/0.95 NA water-immersion objective and an external non-descanned detector, at an acquisition rate of 40 frames/s. VIP-IN Ca²⁺ transients were evoked by electrical stimulation in SLM using a bipolar theta-glass microelectrode and a brief train stimulation protocol (100 µA, 100 Hz, 1 s). Jaws illumination was delivered using the same optical fiber used for in vivo optogenetic experiments, positioned on the slice surface above the hippocampal region to approximate the in vivo light-delivery configuration. VIP-IN Ca²⁺ responses were recorded before and during 620-nm illumination of Jaws-expressing MnR axons. Calcium transient amplitude was quantified as peak ΔF/F in the response window following SLM stimulation. Responses obtained during Jaws illumination were compared with control stimulation responses recorded in the same cells. Because electrical stimulation in SLM activates mixed local and afferent inputs, this experiment was interpreted as a functional validation that the in vivo optogenetic illumination protocol inhibits a MnR-sensitive component of synaptically evoked VIP-IN recruitment.

Electrophysiological signals were acquired using a Multiclamp 700B amplifier, filtered at 2-3 kHz, digitized at 10kHz with a Digidata 1440 interface, and recorded using the Clampex 10.5 software (Molecular Devices).

### Pharmacology

The following pharmacological agents were applied in the bath: NBQX (12.5 µM, Abcam Biochemicals, Cat #: ab120046), D,L-AP5 (100 µM, Abcam Biochemicals, Cat #: ab120271), serotonin hydrochloride (10 µM, Sigma, Cat #: H-9523), MDL-72222 (100 µM, Tocris, Cat #: 0640), RS-102221 (5 µM, Tocris, Cat #: 1050). MnR-EPSC sensitivity to 5-HT3aR and glutamate receptor antagonists was assessed for each cell by comparing EPSC amplitudes recorded during a 10-min baseline period immediately preceding drug application with those measured during a 10-min window following the 10-min drug wash-in. Cells exhibiting a statistically significant reduction in EPSC amplitude in the presence of MDL-72222 were classified as sensitive. For summary presentation, MnR-EPSCs were normalized to baseline EPSC amplitude recorded in each cell prior to drug application.

To examine how 5-HT2cR activation modulates VIP-IN excitability, we recorded membrane responses to stepwise somatic current injections in the presence of 5-HT, and during co-application of 5-HT with the selective 5-HT2cR antagonist RS-102221. The first AP elicited at rheobase was analyzed for threshold (membrane potential at AP initiation determined from the voltage derivative), amplitude (voltage difference between threshold and AP peak), half-width (AP duration measured at 50% of its amplitude), and AHP amplitude (voltage difference between the AP threshold and the most negative membrane potential reached after AP). Firing frequency was quantified as the AP number elicited during current steps from 20 to 100 pA. Measurements obtained in the presence of 5-HT or 5-HT plus RS-102221were compared to control measurements obtained in ACSF.

### Patch-seq and transcriptomic analysis

Transverse hippocampal slices (thickness, 300 µm) were prepared from VIP-eGFP mice (P15–P25) of either sex as described previously (Luo et al., 2019). VIP-positive O/A interneurons were identified by eGFP expression under blue light (450–490 nm). Experimental tools, including glass capillaries, were autoclaved, and working surfaces cleaned with DNA-OFF (Takara, Cat #: 9036) and RNase Zap (Life Technologies, Cat #: AM9780). The patch-clamp protocol was optimized for high-quality single-cell RNA-seq, following established methods (Fuzik et al., 2016; Cadwell et al., 2016; Luo et al., 2019). Patch pipettes (2–4 MΩ resistance) were filled with ∼1 µl of K^+^-based intracellular solution optimized to maximize RNA recovery by including glycogen (20 µg/ml; ThermoFisher Scientific). Positive pressure (0.1–0.2 ml) was applied to the pipette before contacting the cell membrane. Electrophysiological recordings were performed in current-clamp mode to assess membrane properties. Stable cells were held in whole-cell mode for 15–20 minutes to allow biocytin diffusion. RNA was collected by gently applying suction (0.2–0.3 ml) through the pipette until the cell cytoplasm and nucleus were aspirated, ensuring high RNA yield. Samples with debris contamination were discarded. Collected RNA was transferred to RNase-free PCR tubes containing lysis buffer and RNase inhibitor, then stored at –80 °C. Slices were fixed with 4% PFA and processed for biocytin labeling to recover cell morphology.

Negative controls were generated by aspirating tissue from O/A region, while positive controls included pooling three GFP+ cells in a single pipette. cDNA libraries were constructed using the SMART-Seq v4 kit (Clontech Laboratories, Takara Bio; Mountain View, CA, USA), with first-strand synthesis primed by CDS Primer II A, and template switching performed by the SMART-Seq v4 Oligonucleotide at the 5’ end of the transcript. Amplified and purified cDNA underwent quality validation (Agilent Tapestation 2200) and met the concentration threshold of 150 pg/µl (Cadwell et al., 2016; Luo et al., 2019). cDNA libraries were prepared for Illumina Next Generation sequencing using Nextera XT DNA Library Preparation kits (Illumina Inc., San Diego, CA, USA) and sequenced on the Illumina HiSeq 2500 platform (paired-end, 225 bp insert size). Investigators constructing cDNA libraries were blinded to the experimental conditions (cell vs. negative vs. positive control). As an alternative source of transcriptomic data, we have also analyzed the dataset from the hippocampal CA1 interneuron transcriptomic database (Harris et al., 2018; Fig. 2f) and the Allen Brain Map’s mouse whole cortex and hippocampus SMART-seq (Supplementary Fig. 2d).

### In vivo calcium imaging and optogenetic inhibition

Before experimental manipulations, animals were allowed to recover for 2 weeks after GRIN lens implantation procedure. During the second week of recovery, mice were gradually habituated to handling and miniature microscope attachment while remaining in their home cage, followed by exploration of an open arena. Calcium imaging and optogenetic inhibition were performed using the dual channel nVoke system (Inscopix, Bruker Inc). The microscope was equipped with two LEDs: a 455 ± 8 nm LED for GCaMP7b excitation and a 620 ± 30 nm LED for activation of Jaws. The imaging LED was used at excitation powers up to 2 mW, with 0.1 mW power resolution. The 620-nm LED used for optogenetic inhibition was set to 3 mW. Imaging was performed at a maximum field of view of 1280 × 800 pixels, corresponding to approximately 1050 × 650 μm, at 20 frames/s. The focal plane was adjusted electronically for each mouse within a range of up to 200 µm beneath the lens to maximize the number of visible cells and was kept constant across imaging sessions.

Imaging and optogenetics experiments were performed during open-field exploration, contextual novelty transition, and tail suspension testing. VIP-IN activity was recorded from the dCA1 pyramidal layer, where VIP/CR-IS-3 and VIP/CCK-BC somata are located and where VIP-IN density is highest relative to other CA1 layers (Luo et al., 2020). To test the role of MnR**→**dCA1 projections during novelty, Jaws-mediated optogenetic inhibition (Chuong et al., 2014) was applied in closed loop upon entry into the novel compartment. A continuous 620-nm light pulse was delivered for 90 s beginning when the animal entered the novel context. Control sessions were performed without 620-nm optogenetic illumination. For open-field experiments, control and opto-inhibition epochs were acquired during the same session and in counterbalanced order across mice, as indicated below. For the tail-suspension test, mice were suspended by the tail for 90 s while calcium activity was recorded under control vs. MnR opto-inhibition. Animal behavior was recorded simultaneously using a video camera (Imaging Source USB or Digital USB 2.0 CMOS Camera, 60516, Stoelting Co.) synchronized with the ANY-maze software (Stoelting Co.). Arenas and objects were cleaned with 70% ethanol between animals to minimize odor cues.

### In vivo calcium imaging analysis

Miniscope recordings were processed using Inscopix Data Processing Software. Recordings were preprocessed to remove artifacts, spatially filtered to improve signal-to-noise ratio, and motion-corrected to ensure stable alignment of pixels across frames. Regions of interest and fluorescence traces were extracted using CNMF-E. Extracted calcium traces were exported as .csv files and analyzed using Igor Pro 9.0 and custom Python scripts.

For novelty analysis, extracted calcium traces were aligned to entry into the novel context and converted to median-based Z-scores for visualization and quantitative analyses. Novelty responses were normalized to the median calcium signal during the final 30 s spent in the familiar compartment immediately before entry into the novel compartment. Novelty-evoked activity was quantified during the first minute after entry into the novel environment. A novelty modulation index (NMI) was calculated for each cell as:

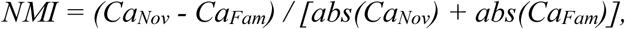

where Ca_Nov_ is the mean calcium signal during the first 60-s novelty exploration and Ca_Fam_ is the mean calcium signal during the last 60-s in familiar environment before door opening. Absolute values in the denominator were used as appropriate because Ca_Nov_ and Ca_Fam_ can be negative. Cells were classified as Nov+, Nov−, or non-responsive according to the sign and significance of their novelty modulation index and the 2SD response threshold.

To quantify speed modulation of VIP-IN activity, animal speed was aligned to calcium traces. Mobility was defined as speed >2 cm/s and immobility as speed ≤2 cm/s. A speed-modulation index (SMI) was calculated as:

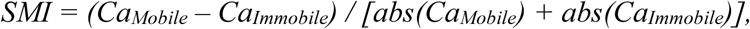

where Ca_Mobile_ and Ca_Immobile_ correspond to the mean calcium signal during mobile and immobile epochs, respectively. Spearman’s rank correlation was also calculated between each cell’s calcium activity and instantaneous running speed. Cells with a significant positive correlation and Spearman’s r ≥ 0.25 were classified as positively velocity-modulated cells (PVMs), whereas cells with a significant negative correlation and Spearman’s r ≤ −0.25 were classified as negatively velocity-modulated cells (NVMs). Cells that did not meet either criterion were classified as non-velocity-modulated (non-VM) cells.

To determine whether novelty-related VIP-IN responses could be explained by increased locomotion, matched-speed analyses were performed. Calcium activity was compared between familiar and novel contexts using epochs within the shared range of running speeds sampled in both conditions, defined as the overlap between the 1st–99th percentile speed distributions of the familiar and novel epochs. This shared speed range was divided into quantile bins computed from the pooled familiar and novel speed values, such that identical bin edges were applied to both conditions. Quantile rather than fixed-width binning was used because running-speed distributions are strongly right-skewed. Within each speed bin, median calcium activity was computed separately for familiar and novel epochs, and the context difference was calculated as Novel – Familiar. Medians were used to limit the influence of large calcium transients. Bins were included only if they contained at least 30 frames from each epoch.

To distinguish an additive novelty-related shift in VIP-IN activity from a change in speed sensitivity, a general linear model (GLM) was fitted to calcium activity sampled within the shared speed range. The model included running speed, acceleration, context, and a context *×* speed interaction term:

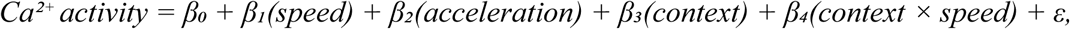

where context was coded as familiar or novel. The context coefficient estimated the speed-independent novelty-related shift in calcium activity, whereas the context *×* speed interaction estimated whether the relationship between speed and calcium activity differed between familiar and novel contexts. Speed and acceleration were included to control for locomotor dynamics. Significance was assessed using block sign-flip permutation testing with 2000 iterations, rather than standard parametric formulas, because consecutive calcium-imaging frames are temporally correlated. Cells with insufficient overlap in running-speed range between familiar and novel epochs, or with too narrow a speed range to separate context and speed-interaction effects, were classified as unresolved and excluded from mechanism assignment.

Because multiple cells were recorded from the same animal, key in vivo optogenetic effects were additionally verified after averaging cell responses within each mouse.

### Behavioral paradigms and behavioral analysis

For all behavioral experiments involving optogenetic manipulation, within-animal comparisons were used whenever possible, such that each mouse served as its own control. For open-field experiments in a familiar environment, mice were first familiarized to the arena. During the test session, behavior and calcium activity were recorded for 10 min, consisting of a 5-min control epoch and a 5-min opto-inhibition epoch. The order of control and opto-inhibition epochs was alternated across mice. Average speed, percentage of mobile time, immobility events, turning angle, rearing frequency, and thigmotaxis index were quantified.

For contextual novelty-transition experiments, two rectangular arenas with identical geometry and dimensions, 36 × 48.5 cm, but distinct sensory cues were used for control and MnR opto-inhibition conditions. The two compartments, referred to as Context A and Context B, were distinguished by different wall and floor textures, color tones, and visual cues, including stickers of different geometrical shapes. During Trial 1, mice with the headstage attached were placed in Context A for 5 min while calcium activity and behavior were recorded. Mice were then returned to their home cage for a 30-min inter-trial interval. During Trial 2, mice were reintroduced to Context A for 5 min, after which the hidden door was opened to allow voluntary entry into Context B. Mice were not forced to enter the second compartment. Once the animal entered Context B, the door was closed. During opto-inhibition sessions, MnR**→**dCA1 projections, 620-nm light was applied for 90 s beginning upon entry into the novel compartment. In a subset of control experiments, circular apparatus (diameter, 60 cm) separated by a hidden door onto two distinct sensory contexts (Tamboli et al., 2024) was used as an additional arena to the rectangular one for comparing VIP-IN activity between familiar and novel contexts.

For the familiar-to-familiar transition control, mice were first allowed to explore two connected compartments with the door open for 10 min, allowing both compartments to become familiar. After a 30-min inter-trial interval in home cage, mice were reintroduced to FamA for 5 min, after which the door was opened to allow voluntary transition into FamB (Supplementary Fig. 8). Once the animal entered FamB, the door was closed, and mice were allowed to explore FamB for 5 min. During opto-inhibition sessions, MnR**→**dCA1 projections were inhibited upon entry into FamB using the same timing as in the novel-context-transition experiment. This control was designed to test transition-related arousal and door-opening effects in the absence of environmental novelty.

For the object-location memory task, mice were allowed to explore two similar LEGO® objects in a rectangular arena, 36 × 29 cm, for 5 min during the sample trial. After a 30-min inter-trial interval, one object was moved to a different location, and mice were returned to the arena for a 5-min test trial. MnR**→**dCA1 opto-inhibition was applied during the full duration of the sample trial. Object exploration time and the number of entries into object zones were scored manually. Object exploration was defined as active investigation directed toward the object, including nose orientation toward the object within 2 cm, but not climbing or incidental passing. Displaced object exploration was calculated as time exploring displaced object / total exploration time x 100%. Object-location memory was quantified using discrimination index (DI):

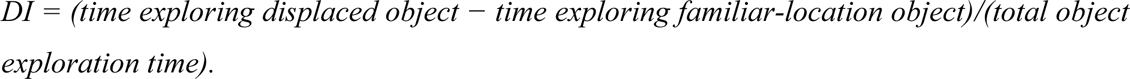

Behavioral data were analyzed using ANY-maze and/or DeepLabCut 3.0.0rc13. DeepLabCut tracking was performed using a top-down HRNet_W32 network architecture initialized by transfer learning from the SuperAnimalTopViewMouse model, with Faster R-CNN used as the detector architecture. For each behavioral video, 20–30 frames were manually labeled using six body keypoints: nose, head, neck, body center, tail base, and tail midpoint. The network was trained for 100 epochs, yielding keypoint likelihood scores of 0.85–0.90. Coordinate data were exported as .csv files, and locomotor metrics were computed using custom Python scripts. Tracking points with low likelihood scores below 0.7 were interpolated before analysis.

Speed was calculated from frame-to-frame displacement of the body center landmark. Mobility was defined as speed >2 cm/s, and immobility as speed ≤2 cm/s. Spatial reorientation was quantified from the animal’s heading direction as the absolute change in movement angle between consecutive trajectory segments. For each session, cumulative absolute turning angle was calculated per second and used as a measure of heading-direction changes during exploration. Rearing events were scored manually and expressed as events per minute. Thigmotaxis was calculated as the proportion of time spent in the peripheral zone of the open field.

For spatial sampling analyses, the novel compartment was divided into 24 spatial bins of 6 × 6 cm. For each bin, the number of entries and total dwell time were quantified. The number of entries per bin was used as a measure of sampling transitions, whereas bin dwell time was used as a measure of local occupancy. Spatial entropy was calculated from the dwell-time probability distribution across the 24 bins:

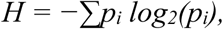

where *p_i_* is the fraction of total binned dwell time spent in bin *i*. Entropy was normalized by the maximum possible entropy, *log_2_(24)*. Spatial-bin measures were analyzed across the full 5-min novel-context exploration period.

For the tail-suspension test, calcium responses were aligned to the onset of suspension and normalized to a baseline period immediately preceding suspension. VIP-IN responses were compared between control and MnR**→**dCA1 opto-inhibition conditions.

### CA1 Microcircuit Model

The computational model was adapted from the biophysically realistic CA1 network model developed by Turi et al. (2019), which included pyramidal cells (PC), axo-axonic cells (AAC), parvalbumin-expressing basket cells (PV-BCs), bistratified cells (BSCs), oriens-lacunosum moleculare cells (OLM), VIP/CCK basket cells (VIP/CCK-BCs), VIP/CR-IS-3 cells, and VIP negative velocity modulated cells (VIP-NVMs). Whereas the original model simulated goal-oriented spatial learning on a one-dimensional linear track, the present version was modified to simulate exploratory behavior in a two-dimensional 100 cm x 100 cm open-field arena during familiar and novel context exploration, with the addition of novelty-modulated MnR inputs to VIP-IN populations.

#### 2D Trajectory and Movement Dynamics

Movement trajectories were simulated in a 100 x 100 cm open-field arena over 22,500 time steps at 1-ms resolution. The spatial position and instantaneous speed of the virtual mouse were generated using a state-dependent stochastic random walk (Bovet and Benhamou, 1988). The simulated animal alternated between active exploration and transient pauses. Heading direction was updated using directional momentum combined with random angular acceleration, and boundary reflections constrained the trajectory within the arena. Instantaneous speed included exponential and Gaussian fluctuations that decayed with friction. To promote exploration of the two-dimensional arena, movement probability and heading direction were also influenced by occupancy history across a 20 x 20 spatial grid, with 5 cm bins. This biased the agent toward less-visited regions of the arena. During the novel-context condition, the probability of movement and the target speed were increased, mimicking the enhanced exploratory engagement observed during novel-context entry.

#### EC, CA3, and MnR Inputs

Spike times were generated from non-homogeneous Poisson processes. Entorhinal cortical (EC) inputs were adapted from Turi et al. (2019) to generate grid-like spatial fields across the two-dimensional arena. As in the original model, grid-like tuning was generated by superimposing inputs of different sizes and phases to produce strong input at defined locations. Two sets of EC inputs were generated, where spike probability was either positively or negatively modulated by speed. Both EC input streams were positively modulated by novelty and by 8-Hz theta phase dynamics. CA3 inputs were generated using a similar approach, but with an additional spatial envelope applied to sharpen position tuning (Lu et al., 2015). In contrast to EC inputs, CA3 inputs were not directly modulated by novelty. MnR inputs were modeled as novelty-modulated inputs to each VIP interneuron subtype: VIP/CR-IS-3, VIP/CCK-BCs and VIP-NVM cells. MnR synaptic conductance values onto different VIP-IN subtypes were based on experimentally estimated values. These values were scaled by a factor of 10 to produce detectable network-level effects in the reduced circuit model.

#### MnR-removal Simulations

To examine the contribution of MnR input to novelty-related circuit dynamics, simulations were performed under control conditions and under MnR-removal conditions. These included global removal of MnR input to modeled CA1 targets, selective removal of MnR input to VIP/CR-IS-3 cells, and combined removal of MnR input to VIP/CCK-BC and VIP-NVM populations. Model output was quantified as the change in firing rate or place-cell density between familiar and novel conditions.

#### Circuit Connectivity

After addition of MnR inputs and VIP-IN circuit interactions, synaptic conductance values were rebalanced to maintain stable network dynamics during two-dimensional familiar-to-novel exploration. Conductance values modified from Turi et al. (2019) are listed in Supplementary Table 2.

#### Place Cell Classification in Simulated Data

Place cells were classified using a custom approach designed for short-duration simulations, in which spatial coverage of the 100 x 100 cm arena was incomplete. Traditional information-theoretic metrics, such as Skaggs spatial information, require more densely sampled occupancy maps and were therefore not used as the primary classification criterion. Instead, simulated pyramidal cells were classified using a sequential procedure based on spike count, local spike density, trajectory-restricted spatial coherence, and comparison with a path-constrained shuffled distribution.

Pyramidal cells with fewer than five spikes were excluded from place cell classification. For the remaining cells, two-dimensional spike coordinates were analyzed using density-based spatial clustering algorithm (DBSCAN implemented in scikit-learn library; Pedregosa et al., 2011; Ester et al., 1996). The neighborhood radius (ε) was set to 20 cm, with a minimum cluster size of five spikes. A cluster was considered a putative place field if its horizontal and vertical bounding dimensions were each within 40 cm (Wilson & McNaughton, 1993), and if the total bounding-box area did not exceed 1600 cm^2^.

To quantify spatially localized firing without penalizing cells for unvisited regions of the arena, trajectory-restricted spatial coherence was computed (Muller & Kubie, 1989). Occupancy and spike-count maps were constructed using 5 cm x 5 cm spatial bins, and firing-rate maps were calculated only for visited bins. Spatial coherence was defined as the Pearson correlation between the firing rate of each visited bin and the mean firing rate of its neighboring visited bins. Because the short simulated trajectory sampled only a narrow subset of the arena, bins were included if they shared at least one visited neighbor, preserving information near the entry and exit points of the trajectory.

To determine whether spatial coherence reflected coordinate-specific tuning rather than temporal spike-train structure (e.g., theta-modulated bursts; O’Keefe and Recce, 1993), a localized permutation test was performed using 100 shuffles. For each iteration, the full spike-time sequence was circularly shifted relative to the trajectory by a random interval between 2000 and 5000 ms (Henrikson et al., 2010). Trajectory-restricted spatial coherence was re-computed for each shuffled iteration to generate a cell-specific null distribution. A pyramidal cell was classified as a place cell if its empirical spatial coherence exceeded the 95th percentile of its own shuffled distribution, provided that it contained a validated DBSCAN cluster or exceeded an absolute baseline coherence threshold.

### Quantification and statistical analysis

Electrophysiological data were analyzed using Clampfit 10.5 and Igor Pro 9.0. Calcium-imaging data were analyzed using Inscopix Data Processing Software, CNMF-E, Igor Pro 9.0, and custom Python scripts. Behavioral data were analyzed using ANY-maze, DeepLabCut, and custom Python scripts.

Normality was assessed using the Shapiro–Wilk test or Kolmogorov–Smirnov test, as appropriate. Normally distributed paired data were analyzed using paired Student’s t-tests. Non-normally distributed paired data were analyzed using Wilcoxon signed-rank tests, whereas non-normally distributed unpaired data were analyzed using Wilcoxon rank-sum tests. Comparisons across more than two groups were performed using one-way Welch ANOVA, repeated-measures ANOVA, Kruskal–Wallis tests, or Friedman tests, as appropriate. Post hoc comparisons were performed using Dunn’s test with Bonferroni correction, Holm correction, or pairwise non-parametric tests, as indicated in Supplementary Table 1. Fractions or proportions were compared using Fisher’s exact test or Fisher–Freeman–Halton exact test. Paired categorical responses were compared using exact McNemar tests. Tests were two-sided unless otherwise stated.

For matched-speed calcium-imaging analyses, a general linear model was used to test whether VIP-IN activity differed between familiar and novel contexts after controlling for running speed and acceleration. The model included speed, acceleration, context, and a context × speed interaction term. Statistical significance for model coefficients was assessed using block sign-flip permutation testing with 2000 iterations to account for temporal correlations in calcium-imaging data.

For computational-model simulations, paired simulation runs were compared across control and MnR-input removal conditions using paired Student’s t-tests or Wilcoxon signed-rank tests, depending on data distribution. Place-cell classification in simulated data was performed using spatial clustering, trajectory-restricted spatial coherence, and circular-shift permutation testing as described above.

For imaging analyses, cells were treated as individual observations when cell-level responses were analyzed, and the number of contributing mice is reported in the figure legends and Supplementary Table 1. When the same cells, animals, or simulation seeds were compared across conditions, paired tests were used. When groups consisted of different cells, animals, or sections, unpaired tests were used. The definition of n is provided in each figure legend and in Supplementary Table 1 and refers to the number of cells unless otherwise stated. Full statistical details for each figure panel, including n values, unit of analysis, test used, paired or unpaired design, test statistics, correction procedures, and exact p values, are provided in Supplementary Table 1.

Summary data are presented as bar graphs, violin plots, or box plots, with individual data points and paired lines shown when applicable. Box plots show the median as a horizontal line; whiskers are defined according to Tukey’s method. Mean values are indicated by black symbols for control conditions and red symbols for opto-inhibition conditions when applicable. Statistical significance was defined as p < 0.05, with *p < 0.05, **p < 0.01, and ***p < 0.001.

## Data and Code Availability

Patch-Seq transcriptomic data generated in this study have been deposited in the Gene Expression Omnibus (GEO) under accession number GSE294910. Additional datasets supporting the findings of this study, including anatomical, electrophysiological, calcium-imaging and behavioral data, are available from the corresponding author upon reasonable request.

Custom Python/Jupyter notebooks used for calcium-imaging and behavioral analyses are publicly available at GitHub: https://github.com/lstopolnik/Calcium-Data-Analysis.

The computational model code used for the reduced CA1 familiar-to-novelty simulations, including control simulations, global MnR-input removal, pathway-specific MnR-input removal, and place-cell analysis, is publicly available at GitHub: https://github.com/agmccrei/Turi_Model_MnR_Simulations.

The model code is adapted from the CA1 circuit model simulations reported in Turi et al. (2019).

## Acknowledgements

We thank Sarah Côté for technical assistance, and Parisa Iloun and Mohamed El Amine Barkat for help with tissue preparation from VGLUT3-Cre and SERT-Cre mice.

## Funding

This work was supported by the Canadian Institutes of Health Research (CIHR Grant #MOP-137072, MOP-142447, PJT-165878, PJT-183687) and the Natural Sciences and Engineering Council of Canada (NSERC Grant #342292-2012, RGPIN-2020-04939) to L.T.

## Notes

### Competing Interest Statement

The authors have declared no competing interest.

