## Supplemental data for "Median raphe input to dorsal CA1 shapes VIP interneuron recruitment and novelty-guided spatial memory"

### Supplementary Data:

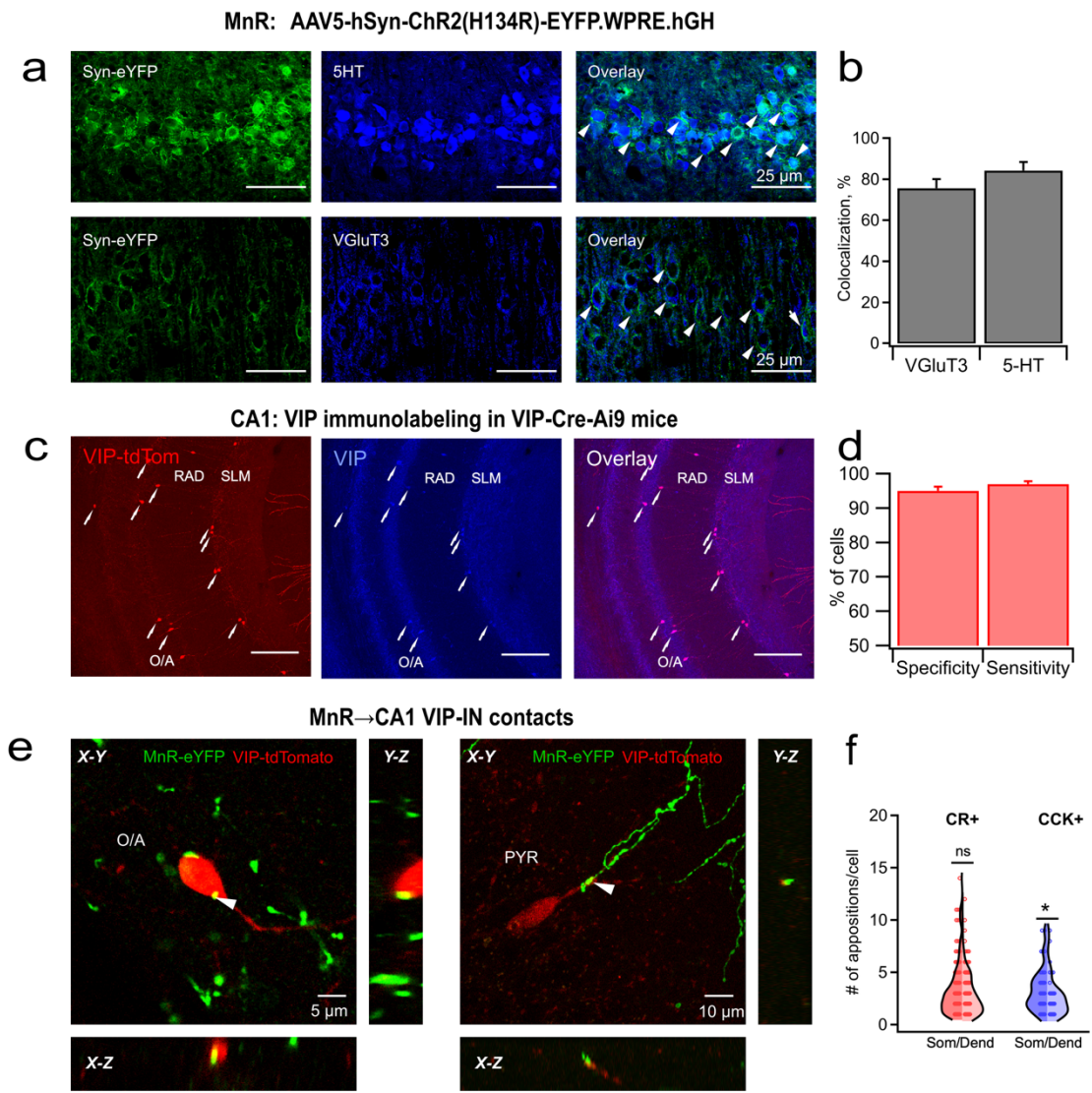

**Supplementary Figure 1. Viral targeting and anatomical validation of MnR inputs to CA1 VIP-INS.** **a,b**, Representative MnR labeling and quantification of eYFP colocalization with 5-HT or VGLUT3 following AAV5-hSyn-ChR2(H134R)-eYFP injection (15 VGLUT3 and 14 5-HT sections from 3 mice, 2 male/1 female). **c,d**, VIP immunolabeling in VIP-IRES-Cre-Ai9 mice and reporter specificity and sensitivity. **e**, Orthogonal confocal views illustrating the criteria for putative somatic and dendritic appositions. **f**, Putative somatic and dendritic appositions onto CR+ and CCK+ VIP-INS (172 CR+ and 45 CCK+ cells from 17 sections and 3 mice). Full statistical details are provided in Supplementary Table 1. \*$p < 0.05$; ns, not significant.

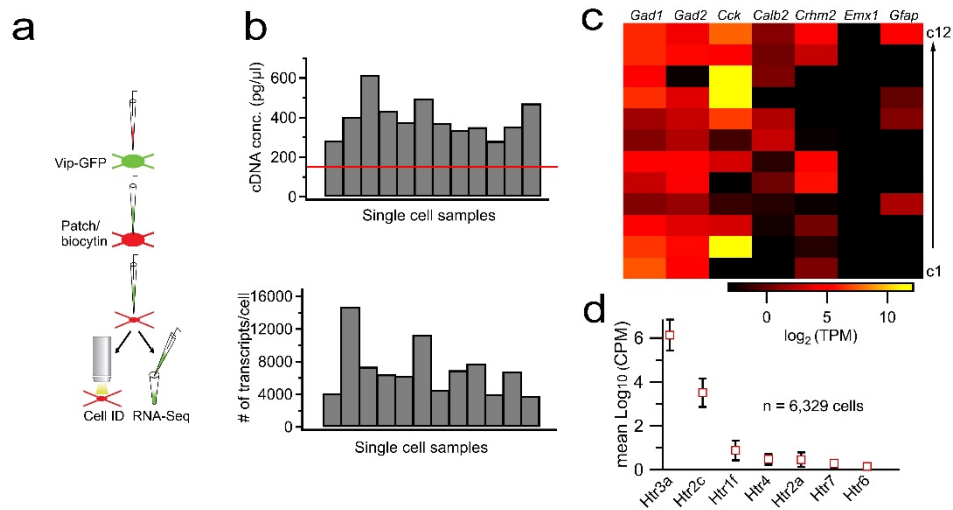

**Supplementary Figure 2. Patch-seq validation of CA1 VIP-IN identity and serotonin-receptor expression.** **a**, Patch-seq workflow. **b**, cDNA concentration and number of detected transcripts for each sample; the red line indicates the 150 pg/μl inclusion threshold. **c**, Expression of interneuron and VIP-IN markers and absence of excitatory-cell marker *Emx1* in 12 cells. **d**, Serotonin-receptor expression across VIP-IN classes in the Allen Brain Map mouse whole-cortex and hippocampus SMART-seq dataset (n = 6,329 cells).

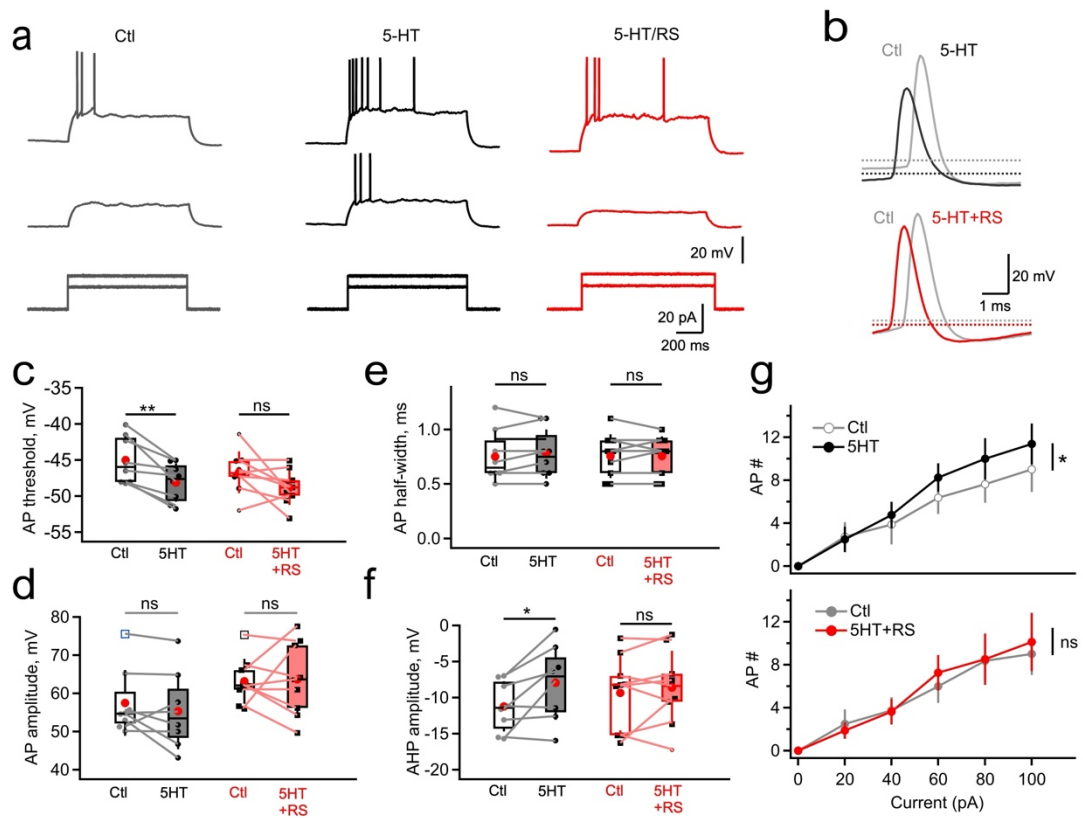

### Supplementary Figure 3. 5-HT<sub>2C</sub> receptor-sensitive modulation of CA1 VIP-IN excitability.

**a,b**, Representative voltage and action-potential traces under control, 5-HT, and 5-HT plus RS-102221 conditions. **c–f**, Action-potential threshold (**c**), amplitude (**d**), half-width (**e**), and afterhyperpolarization amplitude (**f**). **g**, Firing across current steps. (5-HT, 8 cells from 3 male mice; 5-HT plus RS-102221, 9 cells from 6 mice, 3 male/3 female). These data show that 5-HT altered VIP-IN excitability and that the measured effects were attenuated or not detected during 5-HT<sub>2C</sub> receptor blockade. Full statistical details are provided in Supplementary Table 1. \*$p < 0.05$, \*\*$p < 0.01$; ns, not significant.

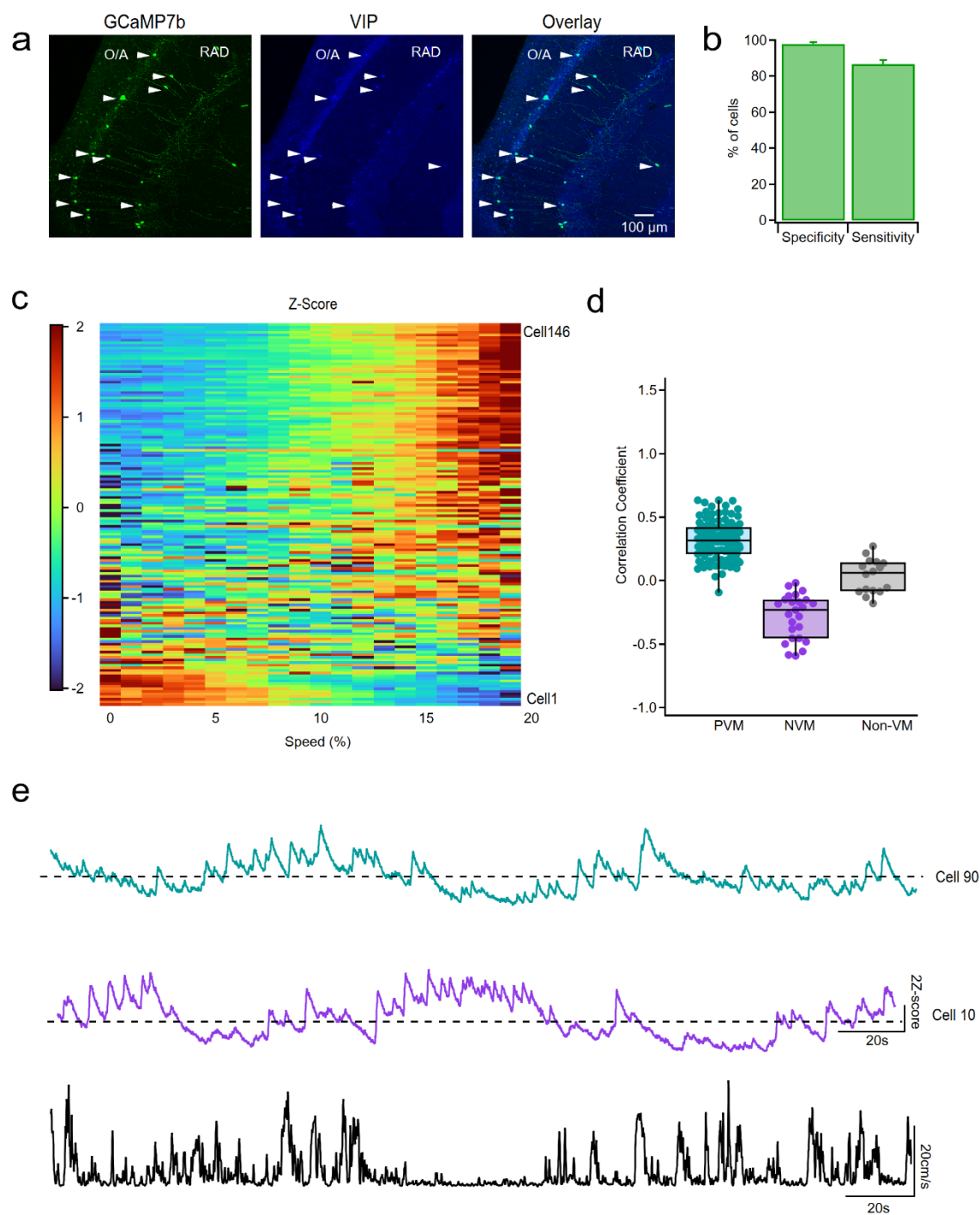

**Supplementary Figure 4. Validation of GCaMP7b expression and velocity modulation in dCA1 VIP-INs.**

**a,b**, GCaMP7b expression, VIP immunolabeling, and targeting specificity and sensitivity. **c**, VIP-IN activity as a function of normalized running speed during familiar exploration ($n = 146$ cells from 11 mice). **d**, Speed–activity correlation coefficients for PVM, NVM, and non-VM classifications. **e**, Representative PVM and NVM activity traces with running speed.

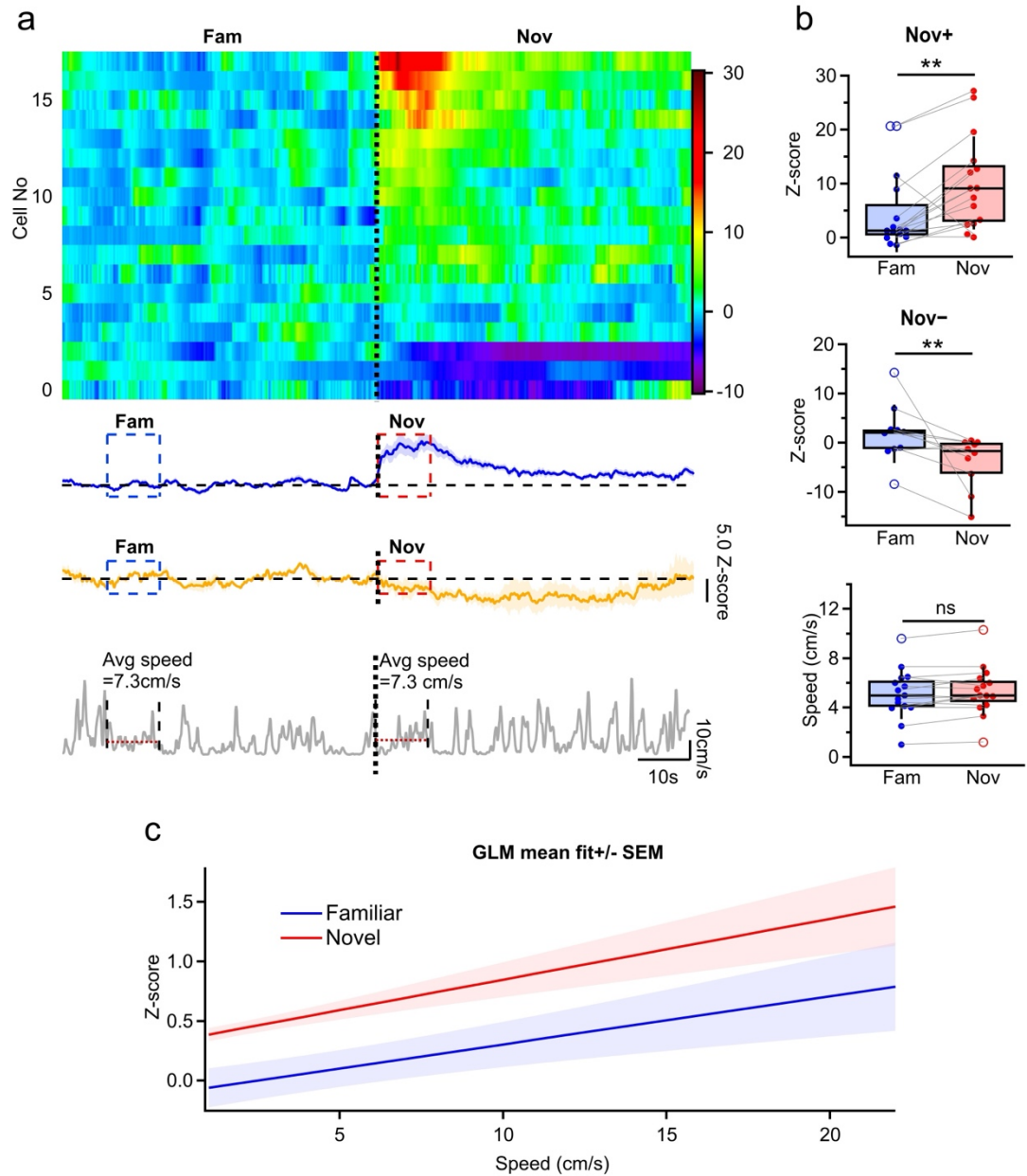

**Supplementary Figure 5. VIP-IN activity in familiar and novel environments at matched running speeds.**

**a**, Representative matched-speed example showing VIP-IN activity from one mouse during familiar and novel epochs. Heat map shows cells from the same imaging session; mean traces for Nov+ and Nov- cells and running speed are shown below. **b**, Animal-wise summary of VIP-IN activity, confirming increased Nov+ VIP-IN activity and decreased Nov- activity during the novel epoch despite matched running speed (15 mice, 6 male/9 female). **c**, Population level general linear model (GLM) analysis showing higher VIP-IN activity in the novel context across the shared speed range after controlling for speed and acceleration. These analyses test whether novelty-associated activity persists at comparable running speeds. Full statistical details are provided in Supplementary Table 1. $**p < 0.01$; ns, not significant.

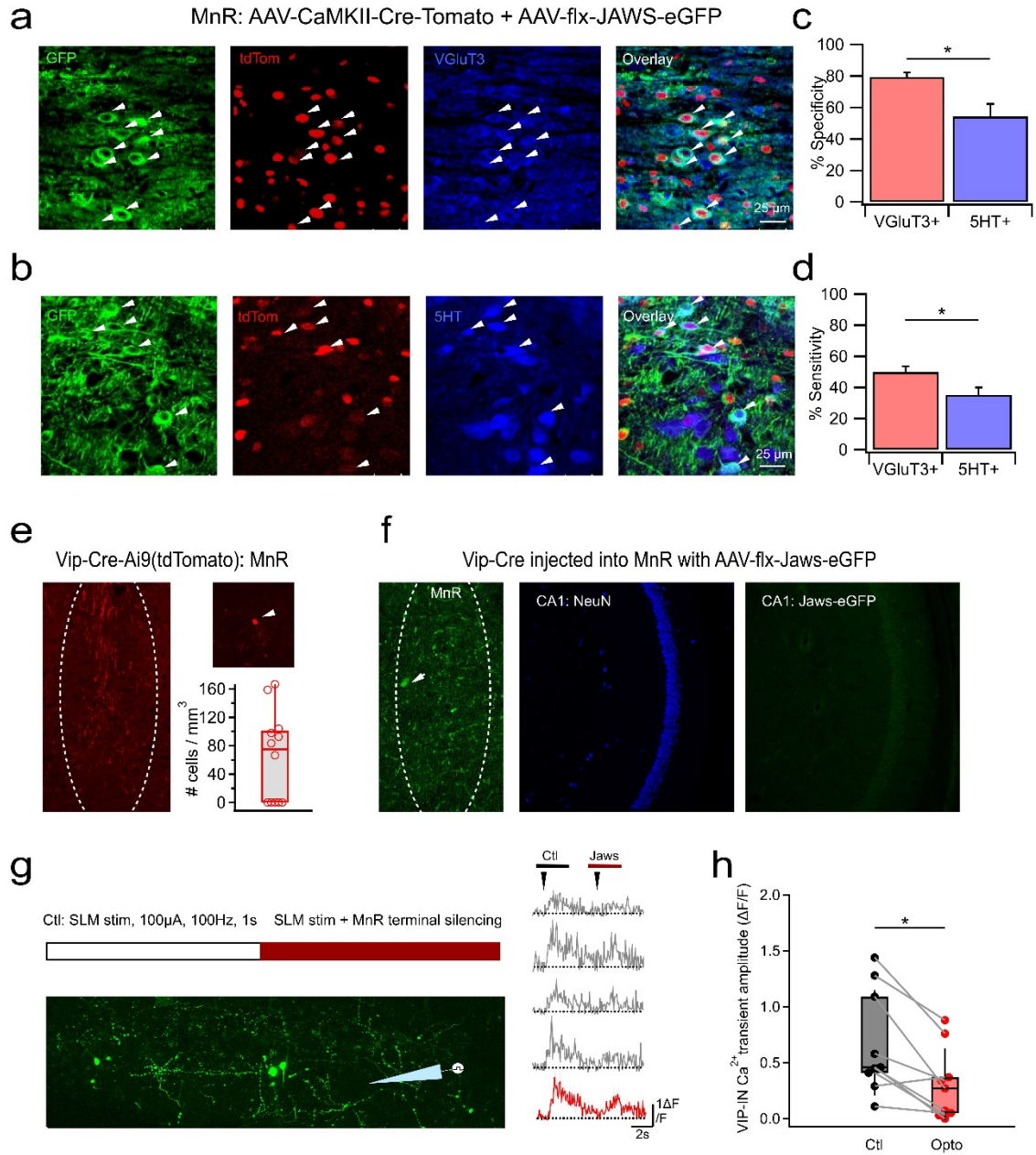

### Supplementary Figure 6. Validation of MnR targeting and Jaws-mediated pathway inhibition.

**a–d**, Jaws expression in VGLUT3+ and 5-HT+ MnR neurons and targeting specificity and sensitivity (33 sections from 3 mice, 2 male/1 female). **e**, Sparse VIP+ neurons and local processes in the MnR. **f**, Lack of detectable CA1-projecting Jaws-eGFP labeling after Cre-dependent viral injection into the MnR of VIP-IRES-Cre mice. **g**, Ex vivo validation protocol and representative VIP-IN calcium responses before and during Jaws illumination. **h**, Jaws illumination reduced the MnR-sensitive component of SLM stimulation-evoked VIP-IN calcium responses ($p = 0.0117$, Wilcoxon rank test; $n = 9$ cells). Full statistical details are provided in Supplementary Table 1. $*p < 0.05$.

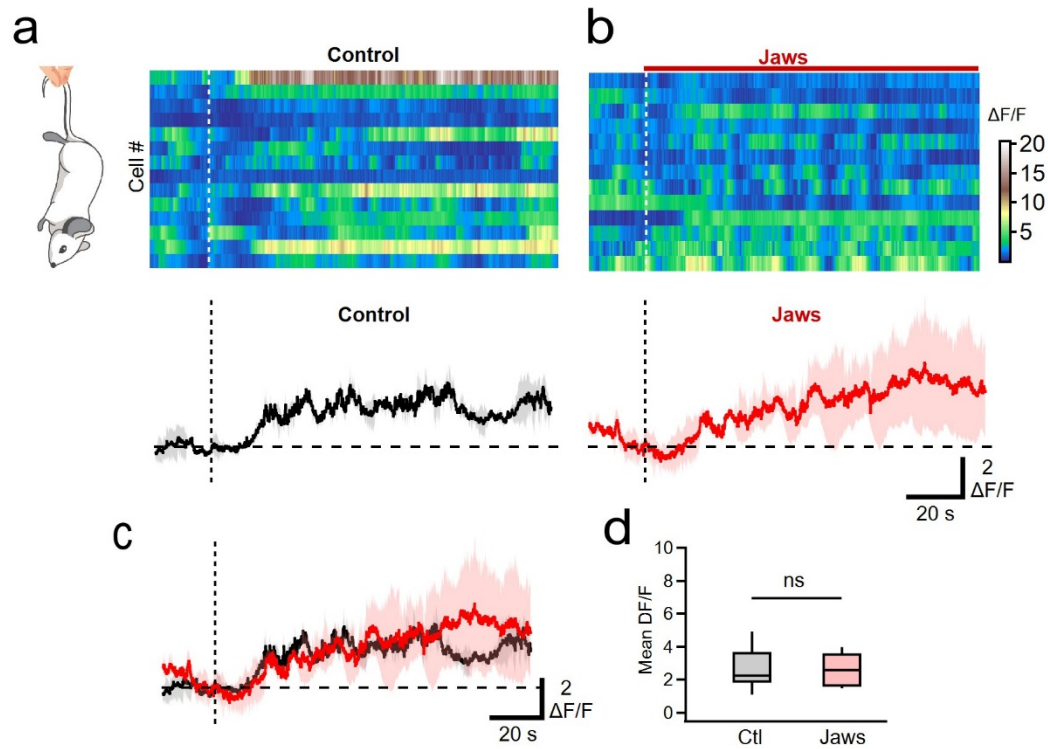

**Supplementary Figure 7. MnR→dCA1 inhibition does not detectably alter VIP-IN activation during tail suspension.**

**a,b,** VIP-IN activity aligned to tail-suspension onset under control and MnR→dCA1 opto-inhibition. **c,** Mean responses across conditions. **d,** Paired comparison of tail-suspension-evoked responses ($p = 0.4725$, Wilcoxon rank test; 38 cells from 3 mice, 1 male/2 female). Both conditions showed activation relative to baseline, but the principal between-condition effect was not detected. ns, not significant.

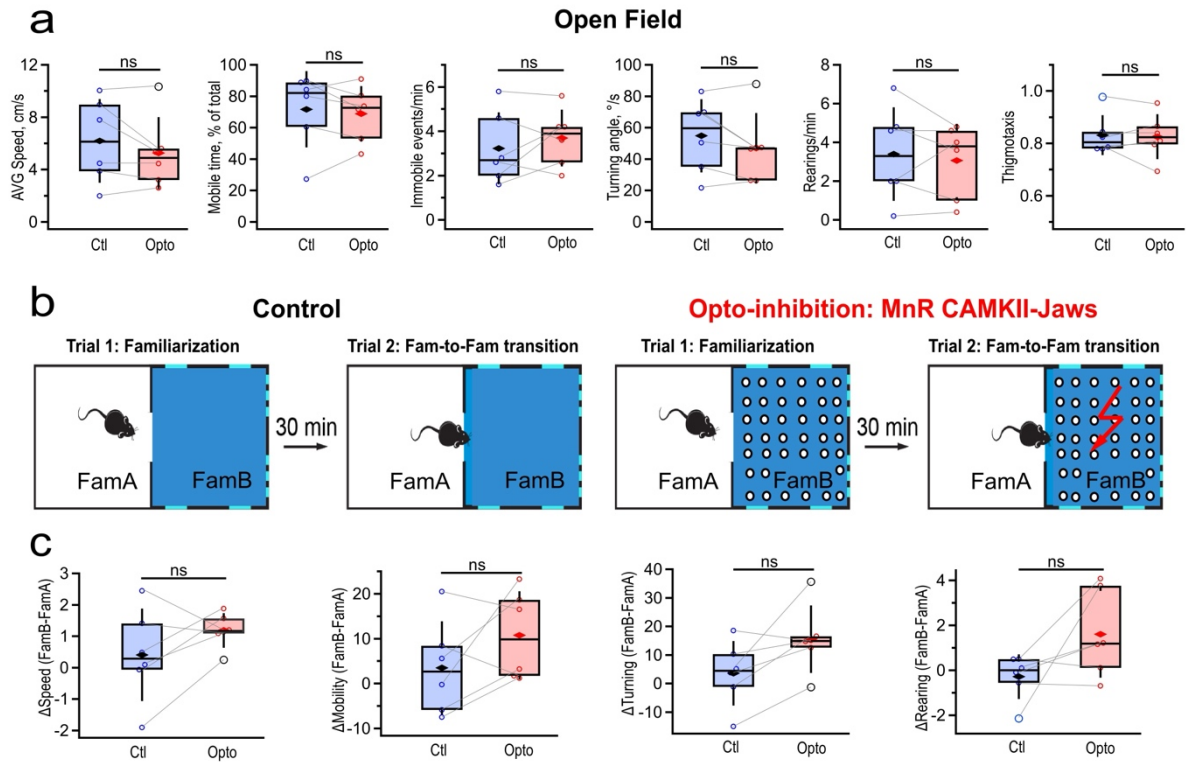

**Supplementary Figure 8. MnR→dCA1 opto-inhibition does not detectably alter familiar open-field behavior or familiar-to-familiar transitions.**

**a**, Average speed, mobile time, immobility events, turning angle, rearing rate, and thigmotaxis during familiar open-field exploration under control and opto-inhibition conditions ($n = 6$ mice, 3 male/ 3 female). **b**, Familiar-to-familiar transition paradigm. **c**, Changes (FamB - FamA) in speed, mobile time, turning angle, and rearing from FamA to FamB ($n = 6$ mice, 3 male/ 3 female). These controls found no detectable broad effect on familiar-environment exploration or movement between familiar compartments. Full statistical details are provided in Supplementary Table 1. ns, not significant.

**Supplementary Table 1. Summary of statistical analysis conducted throughout the study.**

*Note: Animals of both sexes were used throughout the study, with sex distribution balanced across groups whenever possible. Animal sex is indicated in Figure legends.*

| Figure/<br>panel | Measure | n/unit | Comparison | Statistic/<br>p value | Test/Correctio<br>n |
| --- | --- | --- | --- | --- | --- |
| **Fig. 1c** | Mean intensity of eYFP axonal labeling | 24 sections;<br>4 mice | Across CA1 layers;<br>O/A vs. PYR;<br>O/A vs. RAD;<br>O/A vs. SLM;<br>PYR vs. RAD;<br>PYR vs. SLM;<br>RAD vs. SLM | H(3) = 49.24,<br>p = $1.16 \times 10^{-10}$;<br>p = 0.0373;<br>p = 0.9189;<br>p = $3.48 \times 10^{-4}$;<br>p = 0.0817;<br>p = $8.48 \times 10^{-11}$;<br>p = $3.17 \times 10^{-6}$ | Kruskal-Wallis;<br>Dunn/Bonferroni |
| **Fig. 1g** | Distribution of aggregate putative MnR appositions | 172 CR+ cells;<br>45 CCK+ cells;<br>17 sections;<br>3 mice | CR+ vs. CCK+;<br><u>Layer effect:</u><br>CR+ cells;<br>CCK+ cells | H(7) = 27.29,<br>p = 0.0003<br>H(3) = 23.23,<br>p = 0.000036<br>H(3) = 4.08,<br>p = 0.253 | Kruskal-Wallis test<br>Kruskal-Wallis/Holm test |
| **Fig. 1l** | Connection probability out of total recorded | 10/20 IS-3s, 9 mice;<br>8/11 VIP/CCK-BCs, 6 mice;<br>7/10 VIP-OAs, 5 mice | Across VIP-IN classes;<br>IS-3 vs. VIP/CCK-BC;<br>IS-3 vs. VIP-OA;<br>VIP/CCK-BC vs. VIP-OA | p = 0.418;<br>p = 0.275;<br>p = 0.440;<br>p = 1.0 | two-sided Fisher-Freeman-Halton exact test |
| **Fig. 1m** | MnR-EPSC latency | 10 IS-3s, 9 mice;<br>8 VIP/CCK-BCs, 6 mice;<br>7 VIP-OAs, 5 mice | Across VIP-IN classes;<br>IS-3 vs. VIP-BC;<br>VIP-BC vs. VIP-OA | F(2,22) = 12.33,<br>p = 0.0002;<br>p = 0.0013;<br>p = 0.0002 | one-way Welch ANOVA;<br>Wilcoxon rank-sum |
| **Fig. 1n** | MnR-EPSC amplitude | 10 IS-3s, 9 mice;<br>8 VIP/CCK-BCs, 6 mice;<br>7 VIP-OAs, 5 mice | Across VIP-IN classes;<br>IS-3 vs. VIP-BC;<br>VIP-BC vs. VIP-OA | F(2, 22) = 11.09,<br>p = 0.00026;<br>p = 0.0003;<br>p = 0.0025 | one-way Welch ANOVA;<br>Wilcoxon rank-sum |
| **Fig. 2c** | Number of VGLUT3 and 5-HT appositions | VGLUT3: 86 CR+ and 38 CCK+ cells, 19 sections, 2 mice;<br>SERT: 63 CR+ and 26 CCK+ cells, 19 sections, 3 mice | <u>VGLUT3</u>: CR+ vs. CCK+;<br><u>SERT</u>: CR+ vs. CCK+;<br><u>CR+ cells</u>: VGLUT3 vs. SERT appositions | p = 0.3856;<br>p = 0.7917;<br>p = 0.013 | Wilcoxon rank-sum |
| **Fig. 2g** | Fraction of cells sensitive to drugs | MDL-sensitive: 7/11 cells,<br>AP5+NBQX-sensitive: 11/12 cells, 12 mice | MDL-72222 vs. AP5+NBQX | p = 0.125 | exact two-sided McNemar test |
| **Fig. 2i** | MnR-EPSC amplitude | 7 cells, 6 mice | Ctl vs. MDL;<br>MDL vs. AP5+NBQX;<br>glutamate vs. 5-HT3aR component | p = 0.0156;<br>p = 0.00183;<br>p = 0.0313 | Wilcoxon signed-rank;<br>Wilcoxon rank-sum |
| **Fig. 3h** | Mean activity | Nov+: 109 cells, 11 mice;<br>Nov-: 32 cells, 8 mice | Nov+ F<sub>A</sub> vs. N<sub>1st</sub> vs. N<sub>3rd</sub>;<br>Nov+ F<sub>A</sub> vs. N<sub>1st</sub>;<br>Nov+ F<sub>A</sub> vs. N<sub>3rd</sub>;<br>Nov+ N<sub>1st</sub> vs. N<sub>3rd</sub>;<br>Nov- F<sub>A</sub> vs. N<sub>1st</sub> vs. N<sub>3rd</sub>;<br>Nov- F<sub>A</sub> vs. N<sub>1st</sub>;<br>Nov- F<sub>A</sub> vs. N<sub>3rd</sub>;<br>Nov- N<sub>1st</sub> vs. N<sub>3rd</sub>; | F(2, 216) = 67.938, p = 5.67 x 10<sup>-20</sup>;<br>t(108) = -9.12 p = 1.40 x 10<sup>-19</sup>;<br>t(108) = -2.3582 p = 0.0605;<br>t(108) = 10.43 p = 1.45 x 10<sup>-17</sup>;<br>$\chi^2(2) = 41.312$, p = 1.069 x 10<sup>-9</sup>;<br>p = 1.40 x 10<sup>-9</sup>;<br>p = 2.36 x 10<sup>-7</sup>;<br>p = 0.1074 | rmANOVA<br>paired t-test/Bonferroni;<br>Friedman test;<br>Wilcoxon signed-rank |
| **Fig. 3j** | Speed Modulation Index | Nov+: 109 cells, 11 mice;<br>Nov-: 32 cells, 8 mice | Nov+ vs. Nov- cells | p = 6.68 x 10<sup>-5</sup> | Wilcoxon rank-sum |
| **Fig. 3l** | Fraction of Nov+ vs. Nov- cells within speed groups | PVM: 105 cells, 11 mice;<br>NVM: 25 cells, 8 mice;<br>Non-VM: 16 cells, 6 mice | PVM vs. NVM;<br>NVM vs. Non-VM;<br>PVM vs. Non-VM | p = 4.1 x 10<sup>-5</sup>;<br>p = 0.0124;<br>p = 0.2574 | Two-sided Fisher exact test |
| **Fig. 3m** | Novelty Modulation Index | PVM: 105 cells, 11 mice;<br>NVM: 25 cells, 8 mice;<br>Non-VM: 16 cells, 6 mice | PVM vs. NVM vs. Non-VM;<br>PVM vs. NVM;<br>PVM vs. non-VM;<br>NVM vs. non-VM | H(2) = 18.4726, p = 9.74 x 10<sup>-5</sup>;<br>p = 0.0002;<br>p = 1.0;<br>p = 0.0022 | Kruskal-Wallis test<br>Dunn/Bonferroni |
| **Fig. 4e** | Fraction of Nov+ vs. Nov-/NR cells | Control: 97/114<br>Nov+; MnR-: 74/92 Nov+; 8 mice | Ctl vs. MnR- | p = 0.4561 | Two-sided Fisher exact test |
| **Fig. 4f** | Peak novelty response | 97 Nov+ cells, 54 PVM cells, 8 mice | Nov+ cells: Ctl vs. MnR-<br>PVM cells: Ctl vs. MnR- | Cell-wise: p = 8.59 x 10<sup>-9</sup>;<br>Mouse-wise: p = 0.0156;<br>Cell-wise: p = 0.0036 | Wilcoxon rank-sum |
| **Fig. 4h** | Peak novelty response of Nov- cells | 35 cells, 4 mice | Ctl vs. MnR- | Cell-wise: p = 0.9283 | Wilcoxon rank-sum |
| **Fig. 4j** | Frequency | 18 cells/3 mice | Ctl vs. MnR- | p = 0.4142 | Wilcoxon rank-sum |
| **Fig. 4k** | Peak amplitude | 18 cells/3 mice | Ctl vs. MnR- | p = 0.4598 | Wilcoxon rank-sum |
| **Fig. 4l** | AUC | 18 cells/3 mice | Ctl vs. MnR- | p = 0.6727 | Wilcoxon rank-sum |
| **Fig. 5b** | Firing rate change: familiar-to-novel transition | 15 simulations | PCs,<br>PV-BCs,<br>OLMs,<br>VIP/CR-IS-3, | t(14) = 2.23, p = 0.0428;<br>t(14) = 6.88, p = 7.54 x 10<sup>-6</sup>;<br>t(14) = 4.71, p = 0.0003;<br>t(14) = 15.06, p = 4.81 x 10<sup>-10</sup>; | paired t-test;<br>paired t-test;<br>paired t-test;<br>paired t-test; |
|  |  |  | VIP/CCK-BCs,<br>VIP-NVMs | t(14) = 3.83,<br>p = 0.0018;<br>t(14) = 4.33,<br>p = 0.0007 | paired t-test;<br>paired t-test |
| **Fig. 5c** | Firing rate change (novel – familiar) after MnR removal | 15 simulations | <u>VIP/CR-IS-3:</u><br>Ctl vs. All<br>Ctl vs. IS-3<br><u>VIP/CCK-BCs:</u><br>Ctl vs. All<br>Ctl vs. IS-3<br>Ctl vs. CCK/NVM<br><u>VIP-NVM:</u><br>Ctl vs. All<br>Ctl vs. IS-3<br>Ctl vs. CCK/NVM<br><u>OLMs:</u><br>Ctl vs. All<br>Ctl vs. IS-3<br>Ctl vs. CCK/NVM<br><u>PV-BCs:</u><br>Ctl vs. All<br>Ctl vs. IS-3<br>Ctl vs. CCK/NVM<br><u>PCs:</u><br>Ctl vs. All<br>Ctl vs. IS-3<br>Ctl vs. CCK/NVM | t(14) = 40.65,<br>p = $6.2 \times 10^{-16}$;<br>t(14) = 40.65,<br>p = $6.2 \times 10^{-16}$;<br>t(14) = 8.57,<br>p = $6.06 \times 10^{-7}$;<br>p = 0.0215;<br>t(14) = 9.37,<br>p = $2.06 \times 10^{-7}$<br>t(14) = -8.81,<br>p = $4.38 \times 10^{-7}$;<br>t(14) = -22.15,<br>p = $2.68 \times 10^{-12}$;<br>t(14) = 8.75,<br>p = $4.75 \times 10^{-7}$;<br>p = $6.1 \times 10^{-5}$;<br>p = $6.1 \times 10^{-5}$;<br>t(14) = -0.45,<br>p = 0.6611;<br>t(14) = -16.19,<br>p = $1.85 \times 10^{-9}$;<br>p = 0.2293;<br>t(14) = -9.79,<br>p = $1.2 \times 10^{-7}$;<br>t(14) = 18.33,<br>p = $3.5 \times 10^{-11}$;<br>t(14) = 15.08,<br>p = $4.7 \times 10^{-10}$;<br>t(14) = 6.63,<br>p = $1.13 \times 10^{-5}$; | paired t-test;<br>paired t-test;<br>paired t-test;<br>Wilcoxon signed-rank;<br>paired t-test<br>paired t-test;<br>paired t-test;<br>paired t-test<br>Wilcoxon signed-rank;<br>Wilcoxon signed-rank;<br>paired t-test<br>paired t-test;<br>Wilcoxon signed-rank;<br>paired t-test<br>paired t-test;<br>paired t-test;<br>paired t-test |
| **Fig. 5d** | Place cell density change (Nov – Fam) | 15 simulations | Ctl vs. All<br>Ctl vs. IS-3<br>Ctl vs. CCK/NVM | p = 0.0121;<br>p = 0.0107;<br>t(14) = 2.06,<br>p = 0.0582 | Wilcoxon signed-rank;<br>Wilcoxon signed-rank;<br>paired t-test |
| **Fig. 6c** | AVG Speed<br>$\Delta$ Speed (Nov – Fam) | 14 mice | Ctl Fam vs. Nov<br>MnR– Fam vs. Nov<br>Ctl vs. MnR– | p = 0.00012;<br>p = 0.0034<br>p = 0.5016 | Wilcoxon signed-rank |
| **Fig. 6d** | Mobile time | 14 mice | Ctl Fam vs. Nov<br>MnR– Fam vs. Nov | p = 0.00012;<br>p = 0.00012 | Wilcoxon signed-rank |
| | $\Delta$ Mobility (Nov – Fam) | | Ctl vs. MnR– | p = 0.1726 | |
| **Fig. 6e** | Turning angle<br>$\Delta$ Turning (Nov – Fam) | 14 mice | Ctl Fam vs. Nov<br>MnR– Fam vs. Nov<br>Ctl vs. MnR– | p = 0.00012;<br>p = 0.00012<br>p = 0.0031 | Wilcoxon signed-rank |
| **Fig. 6f** | Rearing rate<br>$\Delta$ Rearing (Nov – Fam) | 14 mice | Ctl Fam vs. Nov<br>MnR– Fam vs. Nov<br>Ctl vs. MnR– | p = 0.00024;<br>p = 0.00012<br>p = 0.0419 | Wilcoxon signed-rank |
| **Fig. 6g** | Entries/bin/min in Nov | 14 mice | Ctl vs. MnR– | p = 0.0494 | Wilcoxon signed-rank |
| **Fig. 6h** | Bin dwell time in Nov | 14 mice | Ctl vs. MnR– | p = 0.104 | Wilcoxon signed-rank |
| **Fig. 6i** | Normalized entropy in Nov | 14 mice | Ctl vs. MnR– | p = 0.9153 | Wilcoxon signed-rank |
| **Fig. 6l** | Total object exploration time<br>Displaced object exploration time<br>Discrimination index | 6 mice | Ctl vs. MnR–<br>Ctl vs. MnR–<br>Ctl vs. MnR– | p = 0.6875<br>p = 0.03125<br>p = 0.0156 | Wilcoxon signed-rank |
| **Fig. S1b** | Colocalization of eYFP (viral targeting) with VGLUT3 and 5-HT | VGLUT3: 15 sections;<br>5-HT: 14 sections;<br>3 mice | eYFP+VGLUT3 vs. eYFP+5-HT | p = 0.8649 | Wilcoxon rank-sum |
| **Fig. S1f** | Number of somatic and dendritic oppositions/cell | CR+: 172 cells;<br>CCK+ 45 cells;<br>n = 17 sections;<br>3 mice | CR+: som. vs. dend.<br>CCK+: som. vs. dend. | p = 0.9513<br>p = 0.0249 | Wilcoxon rank-sum |
| **Fig. S3c** | AP threshold | 5-HT: 8 cells/3 mice;<br>5-HT+RS: 9 cells/6 mice | Ctl vs. 5-HT<br>Ctl vs. 5-HT+RS | p = 0.0078;<br>p = 0.0977 | Wilcoxon rank-sum |
| **Fig. S3d** | AP amplitude | 5-HT: 8 cells/3 mice;<br>5-HT+RS: 9 cells/6 mice | Ctl vs. 5-HT<br>Ctl vs. 5-HT+RS | p = 0.1953;<br>p = 0.9102 | Wilcoxon rank-sum |
| **Fig. S3e** | AP half-width | 5-HT: 8 cells/3 mice;<br>5-HT+RS: 9 cells/6 mice | Ctl vs. 5-HT<br>Ctl vs. 5-HT+RS | p = 0.25;<br>p = 0.625 | Wilcoxon rank-sum |
| **Fig. S3f** | AHP amplitude | 5-HT: 8 cells/3 mice;<br>5-HT+RS: 9 cells/6 mice | Ctl vs. 5-HT<br>Ctl vs. 5-HT+RS | p = 0.0391;<br>p = 0.5703 | Wilcoxon rank-sum |
| **Fig. S3g** | Firing rate | 5-HT: 8 cells/3 mice;<br>5-HT+RS: 9 cells/6 mice | Ctl vs. 5-HT<br>Ctl vs. 5-HT+RS | p = 0.039;<br>p = 0.53 | Friedman test |
| **Fig. S5b** | Mean activity at matched speed | 15 mice | Nov+: Fam vs. Nov<br>Nov–: Fam vs. Nov | p = 0.014;<br>p = 0.013 | Wilcoxon signed rank |
| **Fig. S5c** | VIP-IN activity | 37 cells, 4 mice;<br>Frames restricted to shared Fam/Nov speed range | Nov vs. Fam context after controlling for speed and acceleration;<br>Context x speed interaction | $\beta_{\text{context}} = +0.577$;<br>permutation $p = 0.024$;<br>$\beta_{\text{context} \times \text{speed}} = +3.94$;<br>permutation $p = 0.252$ | GLM: activity ∼ speed + acceleration + context + context x speed; block sign-flip permutation, 2000 iterations |
| **Fig. S6c** | CaMKII-viral targeting specificity | 33 sections, 3 mice | VGLUT3 vs. 5-HT | $p = 0.0248$ | Wilcoxon rank-sum |
| **Fig. S6d** | CaMKII-viral targeting sensitivity | 33 sections, 3 mice | VGLUT3 vs. 5-HT | $p = 0.0213$ | Wilcoxon rank-sum |
| **Fig. S6h** | Jaws effect on VIP Ca-transient | 9 cells | Ctl vs. Opto-inhibition | $p = 0.0117$ | Wilcoxon signed rank |
| **Fig. S7d** | Ca-response during tail suspension test | 38 cells, 3 mice | Ctl vs. Opto-inhibition | $p = 0.4725$ | Wilcoxon signed rank |
| **Fig. S8a** | Open-field: AVG Speed; Mobile time; Immobility; Turning angle; Rearing rate; Thigmotaxis | 6 mice | Ctl vs. Opto-inhibition | $p = 0.3125$;<br>$p = 0.5625$;<br>$p = 0.4375$;<br>$p = 0.3125$;<br>$p = 0.6875$;<br>$p = 1.0$ | Wilcoxon signed-rank |
| **Fig. S8c** | Fam-to-Fam difference: AVG Speed; Mobile time; Turning angle; Rearing rate | 6 mice | Ctl vs. Opto-inhibition | $p = 0.2188$;<br>$p = 0.1563$;<br>$p = 0.0625$;<br>$p = 0.1563$ | Wilcoxon signed-rank |

**Supplementary Table 2. Synaptic conductance changes relative to Turi et al. (2019) CA1 circuit model.**

*Note: Conductances not shown remained unchanged from the original model, and all other parameters remain unmodified.*

| Synaptic Connection | Original Weight ($\mu\text{S}$) | Adapted Weight ($\mu\text{S}$) |
| --- | --- | --- |
| EC → PC | 1.60e-4 | 2.00e-4 |
| CA3 → PC | 1.60e-4 | 2.50e-4 |
| EC → AAC | 1.20e-4 | 2.20e-4 |
| EC → PV-BC | 1.00e-5 | 1.20e-5 |
| EC → BSC | 1.50e-4 | 5.00e-4 |
| EC → VIP/CCK-BC | 3.00e-4 | 1.00e-4 |
| EC → VIP/CR-IS-3 | 3.00e-4 | 4.00e-4 |
| EC → VIP NVM | 3.00e-4 | 1.00e-4 |
| CA3 → AAC | 1.20e-4 | 2.20e-4 |
| CA3 → PV-BC | 2.20e-4 | 1.20e-4 |
| CA3 → BSC | 1.50e-4 | 5.00e-4 |
| CA3 → VIP/CCK-BC | 1.05e-4 | 6.00e-4 |
| CA3 → VIP/CR-IS-3 | 1.05e-4 | 1.00e-4 |
| CA3 → VIP-NVM | 1.05e-4 | 4.00e-4 |
| MnR → VIP/CCK-BC | N/A | 1.90e-3 |
| MnR → VIP/CR-IS-3 | N/A | 4.67e-4 |
| MnR → VIP-NVM | N/A | 5.67e-4 |
| PV-BC → PC | 4.00e-4 | 8.00e-4 |
| BSC → PC | 1.02e-3 | 2.20e-4 |
| OLM → PC | 6.00e-4 | 6.00e-3 |
| VIP/CCK-BC → PC | 2.00e-4 | 1.60e-4 |
| PC → BC | 7.00e-4 | 4.00e-4 |
| PC → OLM | 2.00e-4 | 4.50e-3 |
| PC → VIP/CCK-BC | 5.00e-4 | 5.00e-7 |
| PC → VIP/CR-IS-3 | 5.00e-4 | 5.00e-7 |
| PC → VIP-NVM | 5.00e-4 | 5.00e-7 |
| PV-BC → AAC | 1.20e-4 | 1.00e-4 |
| VIP/CCK-BC → PV-BC | N/A | 3.00e-3 |
| VIP/CR-IS-3 → PV-BC | 4.50e-2 | 3.00e-3 |
| VIP-NVM → PV-BC | 4.50e-2 | 3.00e-3 |
| PV-BC → BSC | 2.90e-3 | 2.40e-3 |
| BSC → OLM | 2.00e-5 | 1.00e-5 |
| VIP/CR-IS-3 → OLM | 3.50e-3 | 1.50e-2 |
| VIP-NVM → OLM | 3.50e-3 | 1.00e-3 |
| BSC → VIP/CCK-BC | 8.00e-4 | 1.00e-7 |
| PV-BC → VIP/CCK-BC | 1.20e-3 | 1.00e-7 |
| VIP/CR-IS-3 → VIP/CCK-BC | N/A | 5.00e-3 |
| VIP/CR-IS-3 → VIP-NVM | N/A | 5.00e-3 |
